# Profiling the baseline performance and limits of machine learning models for adaptive immune receptor repertoire classification

**DOI:** 10.1101/2021.05.23.445346

**Authors:** Chakravarthi Kanduri, Milena Pavlović, Lonneke Scheffer, Keshav Motwani, Maria Chernigovskaya, Victor Greiff, Geir K. Sandve

## Abstract

**Background:** Machine learning (ML) methodology development for classification of immune states in adaptive immune receptor repertoires (AIRR) has seen a recent surge of interest. However, so far, there does not exist a systematic evaluation of scenarios where classical ML methods (such as penalized logistic regression) already perform adequately for AIRR classification. This hinders investigative reorientation to those scenarios where further method development of more sophisticated ML approaches may be required.

**Results:** To identify those scenarios where a baseline method is able to perform well for AIRR classification, we generated a collection of synthetic benchmark datasets encompassing a wide range of dataset architecture-associated and immune state-associated sequence pattern (signal) complexity. We trained ≈1300 ML models with varying assumptions regarding immune signal on≈850 datasets with a total of ≈210’000 repertoires containing ≈42 billion TCRβ CDR3 amino acid sequences, thereby surpassing the sample sizes of current state-of-the-art AIRR ML setups by two orders of magnitude. We found that L1-penalized logistic regression achieved high prediction accuracy even when the immune signal occurs only in 1 out of 50’000 AIR sequences.

**Conclusions:** We provide a reference benchmark to guide new AIRR ML classification methodology by: (i) identifying those scenarios characterised by immune signal and dataset complexity, where baseline methods already achieve high prediction accuracy and (ii) facilitating realistic expectations of the performance of AIRR ML models given training dataset properties and assumptions. Our study serves as a template for defining specialized AIRR benchmark datasets for comprehensive benchmarking of AIRR ML methods.

## Background

The adaptive immune system is responsible for mounting a tailored immune response against antigens (viruses, bacteria, cancer, self-antigens). The adaptive immune receptors (AIRs) expressed on the cell surface of T cells and B cells recognize and bind antigens [1]. To cover a broad space of potential antigens, AIRs maintain high diversity throughout an individual’s lifetime by a stochastic process called V(D)J recombination [2–4]. For instance, in humans, the expected number of unique T-cell receptors (TCRs) is between 10^7^ and 10^8^, sampled from a set of > 10^14^ potential TCRs [5]. Upon antigen encounter, adaptive immune cells are activated and proliferate, with all daughter cells inheriting the same antigen-specific AIR sequence (clonal expansion). After clearance of the antigen, a fraction of the activated adaptive immune cells mature to a memory stage constituting long-term protection against antigen-reexposure [6]. Therefore, a snapshot of the adaptive immune receptor repertoire (AIRR) by immune repertoire sequencing captures information on the **current and past immune state** of an individual, where patterns corresponding to a specific antigenic response may be traced [7–11].

Previous studies have shown that identical or similar TCRs (where similarity may be defined by edit distance or sharing of sub-sequence motifs) can be observed in multiple individuals that share a similar disease or phenotype [12–16]. Such evidence has become the basis for many further studies that developed machine learning (ML) methods to predict the immune states of individuals based on antigen-specific signatures recorded in AIRR data [17,18]. Methods that use both classical ML and deep learning continue to emerge [19], with the aim of establishing AIRR-ML models for clinical diagnostics [20–31]. A large majority of published methods use either the nucleotide or the amino acid sequence of the complementarity determining region 3 (CDR3) to search for and learn the antigen-specific patterns since the CDR3 loops of AIRs are known to be key determinants of the antigen-specificity of AIRs [11,32,33]. Some of the published methods [22,27,31] aptly considered AIRR classification as a multiple instance learning problem (MIL) [34] consisting of repertoires as bags, receptors as instances, and immune state-associated receptors as witnesses.

Given the continued rise in the development and application of ML methods for immune state prediction, it is imperative to understand the capabilities and limits of ML methods applied to AIRR datasets. However, profiling the baseline performance of AIRR ML methods requires a large suite of benchmark datasets representing a wide range of variable properties of AIRR datasets and immune signals. Although experimental AIRR datasets with large sample sizes are being generated occasionally (e.g., the recent ImmuneCODE database [35] containing SARS-CoV-2-specific TCR datasets), very few experimental studies have generated repertoire-labeled AIRR datasets at a high-resolution and with a repertoire size>500 [25,35,36]. While the limited availability of experimental datasets is a challenge in itself in establishing a baseline performance of AIRR ML methods, the biggest challenge is the **lack of ground truth** related to immune state-associated immune signals. Here, the immune state-associated immune signal refers to the pattern or information encoded in the AIRR dataset that allows to differentiate between two or more immune states (hereafter referred to as ***immune signal* or *signal*** for brevity). In the case of AIRR classification using CDR3 sequences, there exists little concrete, consensus knowledge on the size, shape, incidence levels, and diversity of immune state-associated patterns [11,22]. In other words, it is not known whether the immune state-associated pattern will be in the form of full CDR3 sequences [25,36,37] or short sub-sequences (k-mers) [11,22,38–40], how big and diverse the pool of immune state-associated patterns is, and how frequently the immune state-associated patterns occur in repertoires characterised by a particular immune state. In the absence of knowledge on ground truth immune signals in experimental datasets, an alternative approach to overcoming these challenges is to use **simulated datasets**, where the dataset and signal properties can be controlled while preserving and reflecting the complexities of experimental datasets to a large extent [41].

In this study, we aimed to establish the baseline performance of AIRR ML models by profiling both the capabilities and limits of the models in predicting the immune state labels of repertoires (**Figure 1.a**). The generation of simulated datasets for such an endeavour should cover a diverse set of challenges representing multiple variable properties of the AIRR datasets, immune state-associated immune signals, and ML model assumptions (hereafter collectively referred to as ***properties of AIRR-ML model training set-up***). To this end, we simulated a large suite of distinct benchmark datasets (n≈850 datasets with a total of ≈210’000 repertoires containing approximately 42 billion TCRβ CDR3 amino acid sequences), in which the dataset and signal properties were varied (**Figure 1.c**). To ensure nativeness of the simulated AIR sequences in terms of positional biases, amino acid usage, and sequence length distributions, we generated repertoires according to a human VDJ recombination model provided by OLGA [42]. We then varied (a) immune signal properties (described below), (b) sample size (number of *examples* available for training), (c) repertoire size (number of sequences in each repertoire), (d) class balance (balance between positive and negative class *examples* in training dataset), and (e) noise in the negative class (signal incidence in negative class). Note that the italicised term *examples* commonly used in ML literature refers to repertoires throughout this manuscript. Also, note that the term positive class refers to those repertoires that contain immune-state associated sequence patterns, whereas the term negative class refers to those repertoires where the signal can occur by chance. The immune signal properties that were varied in the benchmark dataset simulations include (a) ***witness rate*** (the rate at which signal occurs in the positive class *examples*) (Figure 1.b), (b) number of k-mers (also referred to as motifs) constituting the signal, (c) the size of signal motifs, (d) whether the signal motifs are continuous, and (e) distributional shift (difference in the witness rates of training datasets and future datasets met by the trained model). We then trained ≈1300 ML models with varying assumptions and complexity to profile the limits and scalability of AIRR ML models. We found that even a baseline method such as logistic regression performed surprisingly well at a witness rate as low as 0.002%, comparable to a level of difficulty observed in AIR-based disease studies [44]. We also characterized several scenarios with increasing levels of signal complexities, in which logistic regression failed to learn the true signal and thus exhibited poor prediction performance. Overall, our findings shed light on the immune signal complexities and the basic dataset properties that can pose a challenge to baseline ML models, and may represent a frontier where the development of novel methodologies by the AIRR ML community is needed.

**Figure 1:**
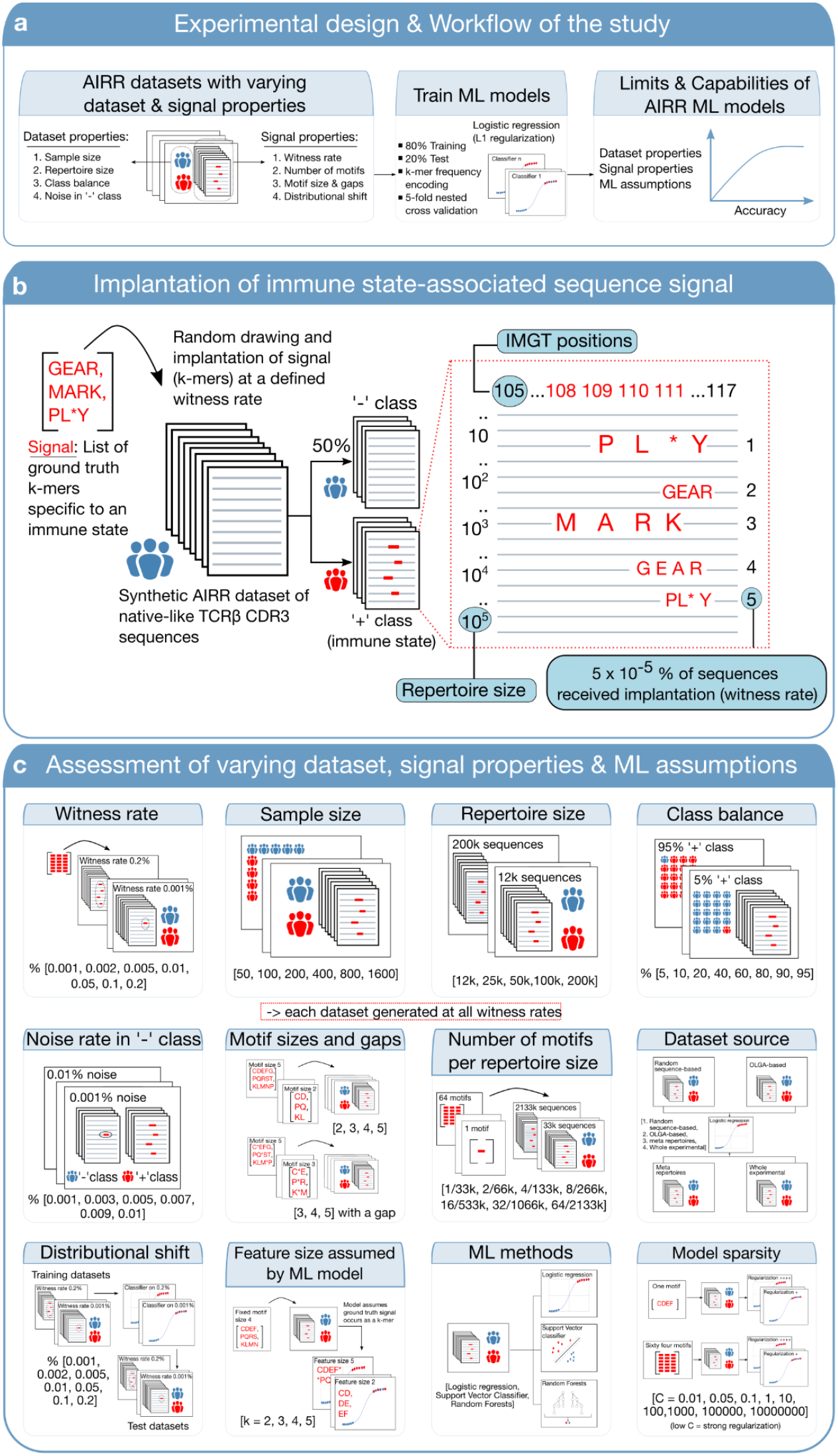
**(a) Experimental design and workflow of the study:** The overarching objective of this study is to profile and establish the baseline limits and capabilities of AIRR ML models. To meet this aim, we evaluate the performance of a baseline logistic regression model across multiple variations of AIRR dataset properties (sample size, repertoire size, class balance, noise in negative class) and immune signal properties (witness rate, number of motifs, motif sizes and gaps, distributional shift). (**b**) **An example of how immune state-associated signal is implanted into AIRR datasets:** To mimic the realistic nature of AIRR datasets, we generate synthetic AIRR reference datasets (specifically TCRβ CDR3 amino acid sequences) according to the VDJ recombination model provided by OLGA [42]. From a list of motifs, we then randomly draw to implant motifs at a defined rate in a defined percentage of repertoires. Note that the wildcard character * in the list of motifs refers to a gap, where any amino acid could occur with equal probability. The repertoires in which motifs are implanted are referred to as positive class and thus contain the patterns associated with a hypothetical immune state. The standard coordinates for CDR3 sequences according to the IMGT numbering system are positions 105-117. Here, IMGT positions [43] refer to the unique numbering system of the ImMunoGeneTics database that positions amino acids in a protein sequence in such a way that facilitates easy comparison of sequences irrespective of the antigen type, chain type, and species. We implant motifs only in the IMGT positions 108-111 with equal probability to not disrupt the positional biases inherent to start and end portions of CDR3 sequences. In this illustration, a total of 5 sequences out of a repertoire size of 10^5^ received a motif implantation. Each positive class repertoire receives a signal at this rate (5 × 10^5^% of sequence; referred to as witness rate throughout this manuscript). (**c) Construction of benchmark datasets with varying dataset and signal properties:** Each signal and dataset property are varied at different rates (shown in square brackets for each property) resulting in multiple separate datasets. For the properties shown within a red-dotted boundary, each separate dataset was further generated at all the witness rates explored (top left). **Table 1** provides a granular overview of the benchmark dataset suite.

### Analyses

The main goal of this study is not to comprehensively assess and benchmark the state-of-the-art machine learning methods for AIRR dataset classification; rather to provide empirical evidence on the baseline performance levels of AIRR ML models across a diverse set of challenges. To this end, we profiled both the capabilities and limits of the models with varying dataset and signal properties (**Figure 1**). Each result is the average of 3 replications of a nested cross-validation in which 80% of the data is used for model training and hyper-parameter tuning, while 20% is used for assessing prediction performance on independent test data. An overview of the variable properties of AIRR ML training setup assessed in this study and their corresponding benchmark datasets is shown in **Table 1, Figure 1.c** and Table S1.

**Table 1:**
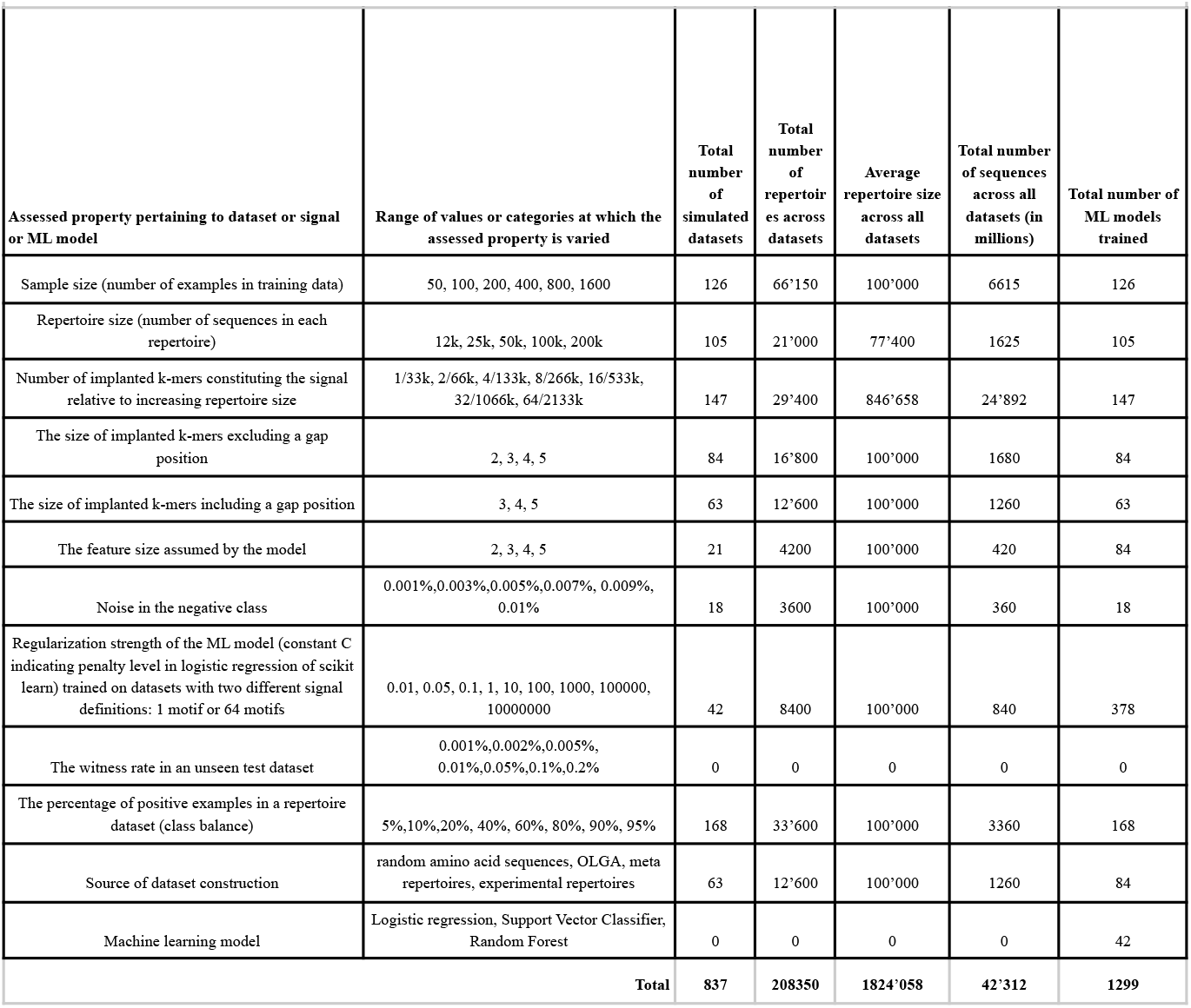
Overview of the variable properties of the AIRR ML training set-up assessed and their corresponding benchmark datasets.

### Impact of k-mer implantation on the background k-mer frequency distributions

Previous studies have shown that the majority of the possible contacts between TCR and peptide antigens were made through only short and typically contiguous stretches of amino acid residues of CDR3s (IMGT positions 107–116) [25,33]. Previously developed ML methods for receptor-specificity (or receptor publicity) prediction or repertoire classification have used such evidence as a premise and often assumed that immune state-associated sequence patterns are short motifs (e.g., 2-mers, 3-mers, 4-mers and 5-mers [20,22,26,45]) while other studies considered the entire AIR sequence [25]. Here, we align with previous observations and profile the baseline performance of AIRR ML models under the assumption that the immune state-associated sequence patterns are short motifs (k-mers). Therefore, to simulate immune state-associated signals in the construction of benchmark datasets, we implanted k-mers into the synthetic AIRR reference datasets (**Figure 1.b**).

We first quantitated the degree to which the implantation of 4-mers affects the background 4-mer frequency distributions of synthetic native-like AIRR datasets (Figure S1; see Methods for details). We focused on 4-mers for this investigation because we used 4-mers as the signal definition in a large majority (≈ 90%) of the benchmark datasets in this study. We observed that when 4-mers were implanted at lower witness rates (up to 10 out of 100’000 sequences receiving implantation), only the implanted 4-mers exhibit significant differences in background frequency distributions. As more sequences received implantation (with an increase of the witness rate), the number of 4-mers that got significantly affected by overlapping partially with the implanted motifs increased. On average, per each implanted 4-mer, a total of ≈35 4-mers that overlapped three residues with implanted motifs, and between 10 to 30 4-mers that overlapped two amino acid residues were significantly disturbed. Very few 4-mers (<four) that were not at all overlapping with the implanted 4-mers exhibited differences in background frequency distributions across three independent replications.

### Impact of model sparsity on prediction performance is dependent on the witness rate and the immune signal definition

The k-mer frequency encoding of AIRR datasets results in high dimensionality. For instance, decomposing a repertoire with 100’000 unique CDR3 sequences into 4-mers would result in approximately 160’000 unique 4-mers (with 20 amino acid residues that can occur at any of the 4 positions of a 4-mer, the total number of possible k-mers is 20^4^ = 160’000). Regularization or shrinkage is useful in high-dimensional problems to avoid overfitting and to improve the generalizability of models. We used Scikit-learn’s [46] implementations of L1 penalized logistic regression models throughout this study, where the hyperparameter controlling regularization strength is indicated by a variable **C**. Smaller values of C represent stronger regularization (see Methods section for details). In order to narrow down an appropriate parameter space for the regularization constant for all the benchmarks in this study, we first evaluated how the prediction performance of models scale with increasing regularization strength. To this end, we explored the impact of regularization strength in two different scenarios with distinct signal definitions, where the defined witness rates are composed of (a) only a single motif and (b) multiple motifs (n=64). At a high witness rate of 0.2%, the signal was so strong that a strong regularization (low C) was not particularly needed to attain higher accuracies (**Figure 2**). This was irrespective of whether the witness rate was composed of a single motif or multiple motifs. As the witness rate decreased, an increase in accuracy was observed with increased regularization strength. When the immune signal was composed of a single motif, a strongly regularized model (C=0.05) was able to classify almost perfectly (99% accuracy) at a witness rate of 0.005% (5 out of 100’000 sequences containing a motif) and performed decently well (82% accuracy) even at a witness rate of 0.002% (2 out of 100’000 sequences containing a motif). Even at a low witness rate of 0.001%, the strongly regularized model performed better than a random prediction by 10 percentage points (**Figure 2.a**). However, when the witness rate was composed of 64 motifs, the models did not exhibit good performance at the lower witness rates (1,2,5,10 sequences out of 100’000 containing a motif), though performed well with a strongly regularized model on witness rates from 0.05% and upwards. For both scenarios (**Figure 2.a** and **Figure 2.b**), the implanted motifs were ascribed higher weights when the models obtained decent performance (Figure S2 and Figure S3). In the case of poor performance, the weights for features corresponding to the implanted motifs were indistinguishable from those of other features (Figure S2 and Figure S3). This confirms that the implanted motifs were indeed the basis for the prediction performance of the models. We further found that the motifs that were ascribed higher weights by the model in this case were the very same motifs that exhibited significant differences in motif frequency distributions as observed through an independent statistical analysis (performed similarly to the previous section) (Figure S4 and Figure S5). This suggests that the users of AIRR classification methods who use a k-mer frequency encoding can use a univariate statistical test either as a feature selection tool or as an additional diagnostic to confirm the basis of classification.

**Figure 2:**
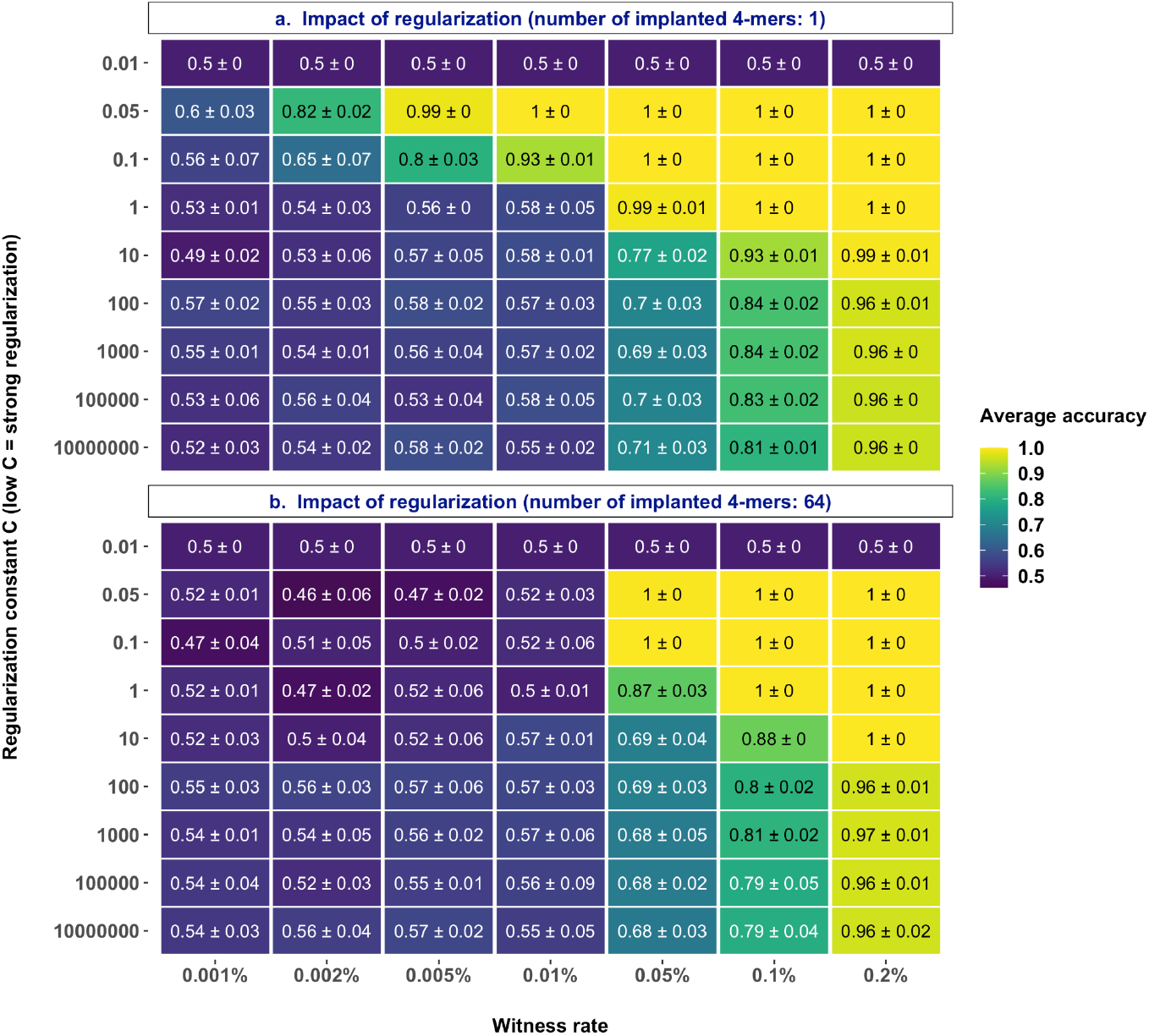
Model sparsity affects prediction performance: Performance estimates of a logistic regression model regularized with a fixed regularization constant C (y-axis) in a binary classification of a balanced, labeled AIRR dataset where the signal in positive class examples composed of 4-mers are known at the explored witness rates (x-axis). The smaller the regularization constant C, the stronger is the regularization. **(a)** Impact of regularization strength when the signal definition is composed of a single 4-mer. (**b)** Impact of regularization strength when the signal definition is composed of 64 4-mers. The mean balanced accuracy of a 5-fold cross-validation was computed in three independent replications. The color coding shows the mean and standard deviation of the performance estimate obtained by three independent replications.

Comparing the frequency distributions of both positive and negative class examples of both scenarios provided additional hints on why a signal definition composed of 64 motifs was a challenging problem to learn for the model (Figure S6, Figure S7). When the signal was composed of 64 motifs, each of the motifs had an equal probability to occur in the sequences of positive class. At a witness rate of 0.002%, in the single motif scenario the same motif will occur in 2 out of 100’000 sequences, whereas in the 64 motifs scenario any two of the 64 motifs will be implanted in each example. This would not only result in less overlap in the signal definition across positive class examples, but also would restrict the implanted motifs from being over-represented relative to baseline frequencies thus making the classes indistinguishable (Figure S7). Such a scenario can arise in experimental studies because of differences in sequencing depth, where the likelihood of detecting more phenotype-associated sequences increases with sequencing depth. These findings indicated that the prediction performance generally improved with model sparsity, though the model struggled at any regularization level as the number of receptors with implanted kmers went below 2 out of the 100’000 receptors for each repertoire.

### Moderate increase in sample size is not sufficient for substantial performance gains

After choosing an appropriate hyperparameter interval for regularization strength (0.05, 0.1, 1, 5), we set out to understand if and how the prediction performance scales with increased sample size (number of repertoires available for training). In the previous section, we observed a decent prediction performance when a single motif was implanted at 0.002% (**Figure 2**). To make the prediction problem slightly more complex, we used a signal definition composed of three motifs in the following experiments unless otherwise stated. This would mean that at 0.002% witness rate, any 2 of the 3 motifs will be implanted independently in each of the positive class examples. To obtain empirical sample complexity estimates, we evaluated the performance with different sample sizes at different witnesses rates (**Figure 3.a**). We observed that at higher witness rates, starting from 0.01% (10 out of 100’000 sequences containing a motif) and above, even a smaller sample size of 50 repertoires was sufficient to reach a decent performance level (78% accuracy). At the lowest witness rate of 0.001%, the explored sample sizes did not provide evidence of a gain in performance even at a sample size of 1600 repertoires. At lower witness rates like 0.002% and 0.005%, there was no substantial gain in performance beyond a sample size of 200 – at a witness rate of 0.002%, the sample size had to increase by 8-fold in order to achieve a performance gain of 5 percentage points. Overall, these findings indicated that a moderate increase in sample size was not sufficient to obtain a substantial gain in the prediction performance. Although there was improvement in prediction performance at lower witness rates, the improvement was moderate and gradual rather than a steep increase.

**Figure 3:**
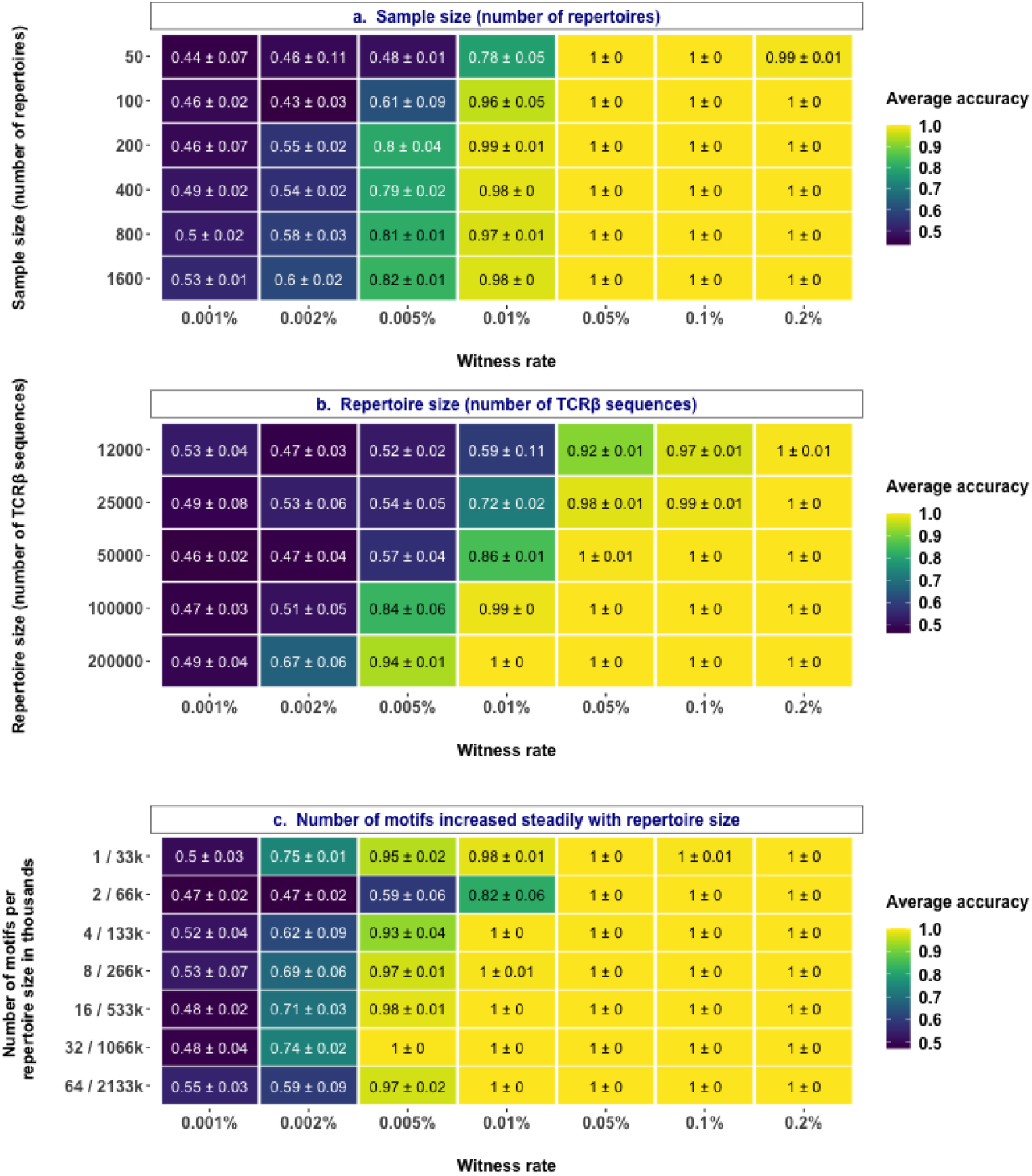
Impact of sample size, repertoire size, and the number of motifs constituting a signal. **(a)** Performance estimates of a regularized logistic regression model in a binary classification of a balanced, labeled AIRR datasets of varying sample sizes (y-axis) where the signal in positive class examples composed of 4-mers are known at the explored witness rates (x-axis). **(b)** Performance estimates of a regularized logistic regression model in a binary classification of a balanced, labeled AIRR datasets with varying repertoire sizes (y-axis) where the signal in positive class examples composed of 4-mers are known at the explored witness rates (x-axis). **(c)** Performance estimates of a regularized logistic regression model in a binary classification of a balanced, labeled AIRR datasets with a combination of varying repertoire sizes and signal definition (y-axis) where the signal in positive class examples composed of 4-mers are known at the explored witness rates (x-axis). The mean balanced accuracy of a 5-fold cross-validation was computed in three independent replications. The color coding shows the mean and standard deviation of the performance estimate obtained by three independent replications.

### Classification performance improves with increase in repertoire size

We then set out to understand if prediction performance would improve with increase in repertoire size (number of TCR sequences per repertoire). For this, we evaluated the prediction performance with different repertoire sizes at different witness rates (**Figure 3.b**). We observed that at higher witness rates, including and beyond 0.05%, the models obtained good performance even at the lowest repertoire size that we explored. At moderate-low witness rates (0.002%, 0.005% and 0.01%), increasing the repertoire size resulted in improved performance by more than 10 percentage points. This was because the increased repertoire size led to a larger absolute count of motifs or sequence patterns associated with the positive class in each repertoire. For instance, at a witness rate of 0.002% in our experiments, a repertoire size of 100’000 would on average carry 2 sequences that contain a phenotype-associated motif whereas a repertoire size of 200’000 would carry 4 sequences that contain the motif. Since the signal definition in this experiment was composed of 3 motifs, it is more likely that a larger portion of signal definition will be included in a repertoire size of 200’000 leading to better performance. Notably, the average number of unique sequences generated in one of the recent experimental studies [25] was around 100’000. Overall, the findings show that prediction performance monotonically increased with increase in repertoire size.

### Impact of repertoire size on performance gain is dependent on the true signal composition

We observed that when the signal definition was composed of many motifs (n=64), all the tested models failed to learn the implanted signal at moderate to low witness rates (**Figure 2**). We also observed that performance gained with increase in repertoire size when the number of motifs was kept constant (n=3) (**Figure 3.b**). Combining the knowledge from both these experiments, we investigated whether and how the performance scales with increased repertoire size if the number of motifs also increases proportionally to the repertoire size. To this end, we trained models at different witness rates on datasets with increasing repertoire size, where the proportion of number of motifs per repertoire size was kept constant. First, we observed that the performance of models gained at moderate to low witness rates (0.002%, 0.005%, 0.01%) even when the number of implanted motifs were 64 (**Figure 3.c**). This is in contrast to the observations that the models performed poorly at moderate to low witness rates when the number of implanted motifs were 64 in repertoires of a fixed size of 100’000 (**Figure 2**). These findings validate the observations that increase in repertoire size contributes to performance gains (**Figure 3.b)**. Irrespective of the increase in the number of implanted motifs, there was a general trend of decent and comparable prediction accuracy at a witness rate of 0.002%, although not necessarily a linear improvement in performance. Overall, the findings validated the performance gains with increased repertoire size but indicated that the impact was dependent on the signal composition (number of implanted motifs) and the relation was non-linear.

### Classification performance strongly depends on class balance in datasets

In the benchmark experiments described above, all the datasets were balanced in labels with 50% each of positive and negative class examples. However, for experimental datasets in biomedical research, it is often a struggle to sample as many negative examples as positive examples. To understand if and how classification performance scales with increased class imbalance, we evaluated the performance at different witness rates with multiple datasets that have varied degrees of class balance. We observed that the class balance did not have a large impact on the performance at moderate to high witness rates (**Figure 4.a**). However, at moderate-low witness rates such as 0.01% and 0.005%, having a high degree of class imbalance (e.g., containing only 20% of examples from either positive or negative class) resulted in a substantial decrease in performance. Overall, these findings demonstrated the dependence of classification performance on class balance.

**Figure 4:**
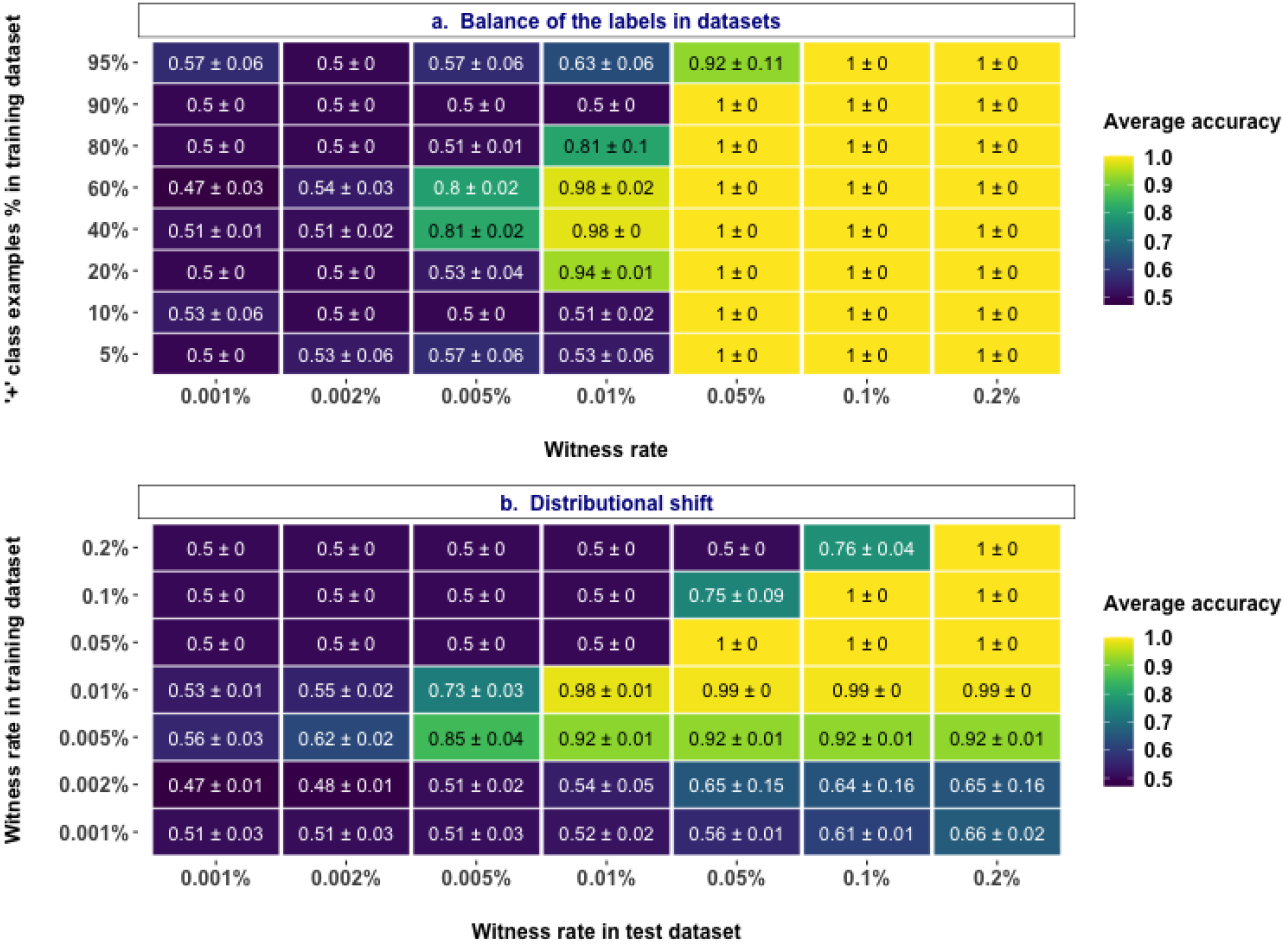
Impact of class balance and distributional shift. **(a)** Performance estimates of a regularized logistic regression model in a binary classification of unbalanced, labeled AIRR datasets with varying degrees of class balance in the datasets (y-axis) where the signal in positive class examples composed of 4-mers are known at the explored witness rates (x-axis). **(b)** Performance of a regularized logistic regression model trained on balanced, labeled AIRR datasets with varying witness rates (y-axis) in the classification of a new unseen balanced, labeled AIRR test datasets where the signal in positive class examples composed of 4-mers are known at the explored witness rates (x-axis). The mean balanced accuracy of a 5-fold cross-validation was computed in three independent replications. The color coding shows the mean and standard deviation of the performance estimate obtained through three independent replicates.

### Distributional shift strongly impacts the classification performance

Distributional shift is a common problem encountered in real world applications of machine learning [47]. It can take many forms but can simplistically be stated as a change in data distributions between training dataset and the examples the model meets in future. To understand how the AIRR ML models that we trained adapt to the distributional shift, we trained the models on datasets with varying witness rates and evaluated their prediction performance on test datasets that have different witness rates (**Figure 4.b**). We observed that the prediction accuracy decreased when the witness rate of either the training or the test dataset decreased. We particularly noticed that the effect of decreased test witness rate was substantial than a decrease in training witness rate. (e.g., a model trained on a dataset with 0.005% witness rate performed better on a test dataset with 0.01% witness rate compared to the performance of a model trained on 0.01% witness rate applied on a test dataset with 0.005% witness rate). Another striking observation was that given a particular training witness rate, a distributional shift that increased the test witness rate improved the performance. Even at very low training witness rates (0.001%), where prediction appears to be essentially random when tested on data from the same distribution, the accuracy increased up to 66% with a distributional shift that increased the test witness rate. This suggests that even though the model trained at 0.001% appears to not having learnt any signal, it has indeed captured the signal, but the signal was not strong enough to allow accurate prediction at its native witness rate; rather if the same signal becomes stronger in an application setting, the same model indeed was able to exploit the signals it has learnt to predict with a decent accuracy.

### Classification performance depends strongly on the shape of the ground truth signal and matched assumptions of the ML model

Previous studies suggested that short and linear sub-sequences (k-mers) of amino acid sequences make contact with the antigenic peptide residues [25,33] and thus many of the previous studies considered the size of ground truth sub-sequences to be between 2–5 amino acid residues that are either continuous or contain gaps [20,22,26,45]. We considered both gapped and ungapped k-mers, which we refer together as ground truth signal shape. We evaluated if and how the performance of models scale with the shape of ground truth signal. To this end, we first assessed the performance of models on datasets where the ground truth had implanted motifs of varied sizes (2-mers, 3-mers, 4-mers and 5-mers) at different witness rates (**Figure 5.a**). We observed that when the signal was longer, a feature size-matched model was able to attain around 65% accuracy even at a witness rate of 0.001% and 80% accuracy at 0.002% witness rate. As the size of signal became shorter, moderate to good performances were observed only at markedly higher witness rates (**Figure 5.a**). Next, we assessed the performance of models on datasets where the ground truth had gapped motifs with varied sizes (3-mers, 4-mers and 5-mers with one gap in random position) implanted at different witness rates. When the true signal contained a gap (e.g., A*C where **\*** could be any single amino acid residue), feature size-matched models did not perform as well as they did when the true signal did not contain any gaps. This was particularly evident at witness rates such as 0.005% and 0.01%, where the performance decreased substantially when compared to a scenario where the true signal did not contain any gaps (**Figure 5.b**). One of the contributory factors for such a decrease in performance might be the fact that the model had to retain multiple coefficients because of the wildcard amino acid residue in the gap position, thus capturing more noise along with the signal.

**Figure 5:**
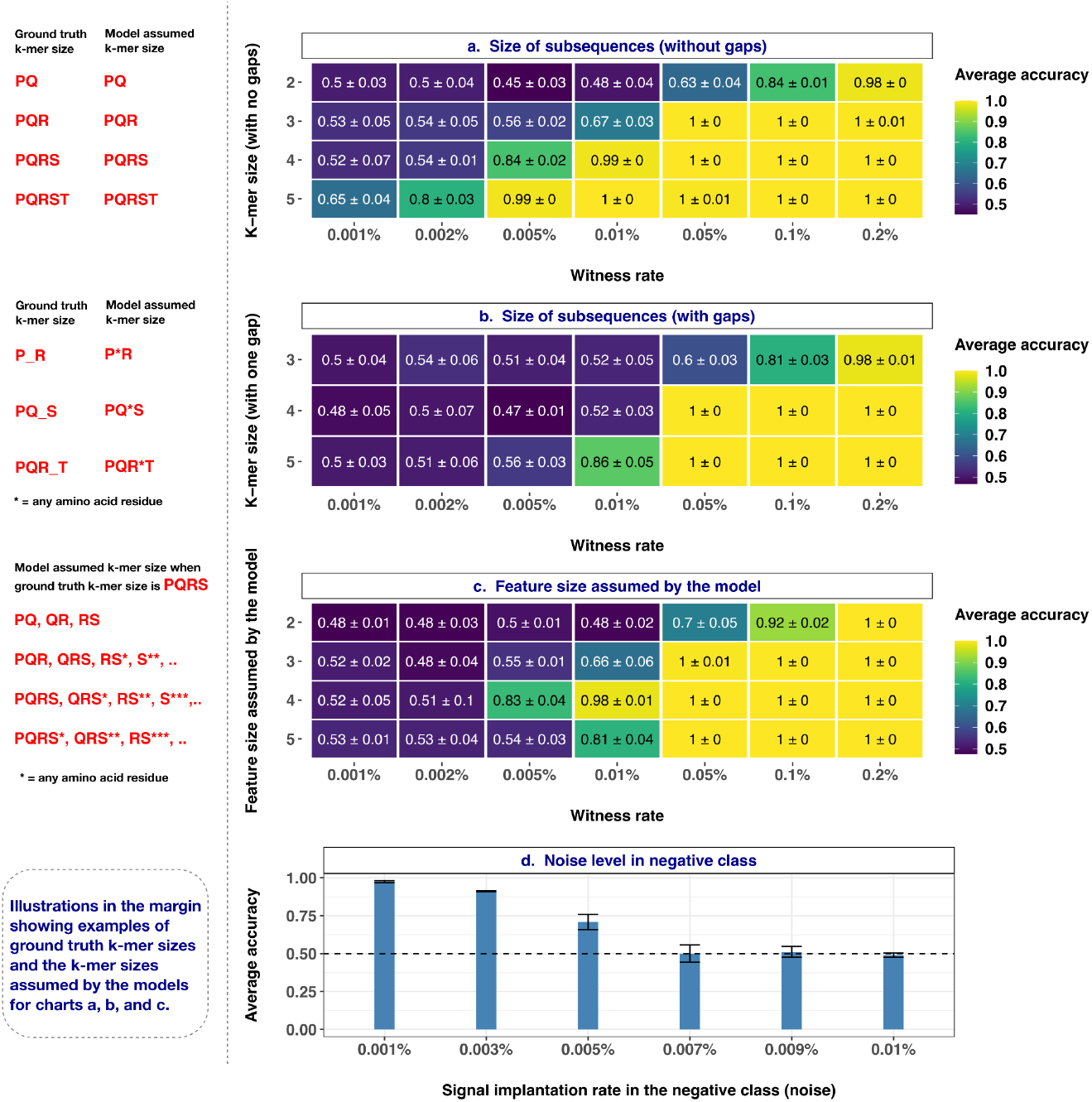
Impact of size of k-mers (with/without gaps), feature size assumed by the model, and noise level in the negative class. **(a)** Performance estimates of a regularized logistic regression model in a binary classification of a balanced, labeled AIRR datasets with the ground truth signal composed of varying k-mer sizes (y-axis) where the signal in positive class examples are known at the explored witness rates (x-axis). **(b)** Performance estimates of a regularized logistic regression model in a binary classification of a balanced, labeled AIRR datasets with the ground truth signal composed of varying k-mer sizes that contained a gap in random position (y-axis), where the signal in positive class examples are known at the explored witness rates (x-axis). **(c)** Performance estimates of a regularized logistic regression model in a binary classification of a balanced, labeled AIRR datasets that assumed varying feature sizes of the ground truth signal (y-axis) where the signal in positive class examples composed of 4-mers are known at the explored witness rates (x-axis). **(d)** Performance estimates of a regularized logistic regression model (y-axis) in a binary classification of a balanced, labeled AIRR datasets where the witness rate in positive class examples composed of 4-mers is fixed at 0.01% and the noise in negative class examples is known at the explored rates (x-axis). The mean balanced accuracy of a 5-fold cross-validation was computed in three independent replications. The color coding shows the mean and standard deviation of the performance estimate obtained by three independent replications. The illustrations in the left margin show examples of ground truth k-mer sizes and the feature sizes assumed by the model for each row of the charts in a, b, c. Note that k-mers shown are only for the ease of illustration; in the simulations a different set of k-mers (n=3) were used.

Further, we set out to understand how the performance of models would vary when the feature size assumed by the model does not match the true signal size. For this, we evaluated the performance of models that made varied assumptions about the size of true signal (assuming the signal occurs as 2-mers or 3-mers or 4-mers or 5-mers) on datasets where the true signal implanted at different witness rates was always held constant as 4-mers (**Figure 5.c**). We observed that when the feature size assumed by the model matches the size of true signal (implanted motif size), the model performed almost perfectly at a witness rate of 0.01%, whereas the performance decreased when the feature size assumed by the model was larger or smaller than the true signal size. A similar trend was also evident at a witness rate of 0.005%. At higher witness rates (>0.01%), a model that assumed a feature size of 2 performed relatively poorly compared to the other models that achieved almost perfect classification accuracy at higher witness rates.

Overall, the findings indicated that learning smaller patterns at lower witness rates was challenging for the tested models compared to learning longer k-mers. The performance at lower witness rates was exacerbated when the ground truth signal contained gaps within the k-mers. ML models with mis-matching assumptions on the ground truth signal fared markedly poorly than the models with matching assumptions.

### Impact of noise on classification performance

In all experiments above, we evaluated the performance of models on synthetic benchmark datasets where we introduced signals into the positive class examples at defined rates. In real-world experiments, one cannot exclude the possibility of various sources of noise in the negative class examples. In this particular ML application, the signal that differentiates positive and negative class examples could for instance occur in both positive and negative class examples but enriched above a baseline in positive class examples. In such a scenario, the background frequencies of the signal in negative class represent noise. To understand the relation between classification performance and the noise levels in negative class, we evaluated the performance of models that are trained on datasets with varying levels of noise (x-axis on **Figure 5.d**) while holding the witness rate in positive class constant at 0.01%. We observed that the level of noise in negative class examples clearly had an impact on the performance of the model. The performance decreased as the noise in the negative class increased. However, the model performed well even when the noise in the negative class was half the witness rate in the positive class.

### Immune receptor sequence-specific biases makes the classification problem unique

Adaptive immune receptor sequences are known to have positional biases, where specific amino acid residues occur with high probability in particular positions of the amino acid sequences. In addition, some amino acid residues might be over-represented in the repertoires and there might be inter-dependencies between the positional frequencies of amino acids. These biases are specific to AIRR datasets. To understand if and how the performance of the tested models differ based on the presence/absence of AIRR dataset-specific biases, we trained and evaluated models on datasets originating from three sources at different witness rates. For this, we generated four different types of datasets as described in the methods section: random sequence repertoires, OLGA-based repertoires, meta repertoires and experimental repertoires. The random sequence repertoires are free of any immune receptor-specific biases and can be a proxy for a special case of general sequence pattern classification problem, while the OLGA-based repertoires contain immune receptor sequences according to the VDJ recombination model of OLGA and are thought to serve as a decent proxy for experimental AIRR sequence datasets. The meta repertoire dataset contains TCRβ CDR3 sequences drawn randomly from repertoires of an experimental dataset irrespective of their immune states or metadata, resulting in repertoires composed of randomly pooled experimentally determined sequences. The experimental repertoires are randomly selected whole experimental repertoires irrespective of their immune states or metadata subsampled to a comparable repertoire size. The findings showed that the performance on random repertoires is poorer at witness rates such as 0.005% and 0.01% when compared to the other two dataset sources (**Figure S8**). Although the performance estimates on OLGA-based repertoires looked similar to those on meta repertoires and subsampled experimental repertoires at 0.005% witness rate, the models on meta repertoires and experimental repertoires exhibited better performance at 0.002%. This points towards the positional biases and dependency structures specific to AIRR datasets.

### Comparison of the performance of selected machine learning methods on AIRR classification

Although a comprehensive assessment and benchmarking of ML methods for AIRR classification is not the goal of this study, we briefly explored how straightforward application of other classical ML methods like Random Forests (RF) or Support Vector Classifier (SVC) compared to the logistic regression model tested throughout this manuscript. To this end, similarly to the exploration of hyperparameter space of logistic regression models, we first explored the hyperparameter spaces of both RF and SVC to narrow down the hyperparameter search space (Figure S9.**b**, Figure S9.c). We then assessed the performance estimates of logistic regression, RF, and SVC in classifying the AIRR datasets with different witness rates. We observed that both RF and SVC exhibited similar performance on this classification problem, but performed poorly compared to the logistic regression model (Figure S9.**a**). We believe this may be due to the stronger degree of regularization of the L1-penalized logistic regression as compared to the other models. Since the goal of this study was not to optimize the models, we did not test other encoding schemes or normalization methods. Nevertheless, the finding points towards the careful need to tailor suitable representations and preprocessing steps and match the ML method assumptions in order to achieve a satisfactory generalizability for the AIRR ML models.

## Discussion

In this study, we observed that a baseline method in the form of a strongly regularized logistic regression model was able to perform well in predicting the immune states of AIRR datasets even at a witness rate as low as 0.002% (2 out of 100’000 sequences containing a motif), comparable to a level of difficulty observed in AIR-based disease studies [44]. We also identified several scenarios where the performance of the baseline method decreased with increase in signal complexities (**Figure 6**). These observations spark curiosity on how well the existing AIRR ML methods would perform on the large and diverse benchmark dataset suite generated in this study. The observation that a baseline method like logistic regression performs exceedingly well in a repertoire classification problem (albeit with a simple definition of potential immune signal) points towards the necessity of carrying out a comprehensive benchmarking of the existing AIRR ML methods for repertoire classification (e.g., see references [22,25,27,28,31,36,37]) to understand if and where tailored ML methods with/without sophisticated architectures are particularly needed to improve the baseline performance. Although benchmarking of existing AIRR ML methods was out of the scope of this study, an observation in a single scenario that logistic regression outperformed two other traditional ML methods (Figure S9) asserts the necessity of benchmarking state-of-the-art AIRR ML methods. To avoid the biases of self-assessment trap [48] (e.g., less transparency in crucial aspects of ML models such as overfitting and poor generalizability) and ensure rigorous assessment of methods in biomedical research, a community-driven concerted effort known as “crowdsourced challenges’’ is increasingly becoming a popular way to benchmark methods [49–51]. The large benchmark dataset suite generated in this study covering a wide variety of properties related to AIRR datasets and immune signals can already act as a suitable subset of scenarios to be assessed in such community-driven benchmarking efforts together with even more specialized benchmark datasets.

**Figure 6:**
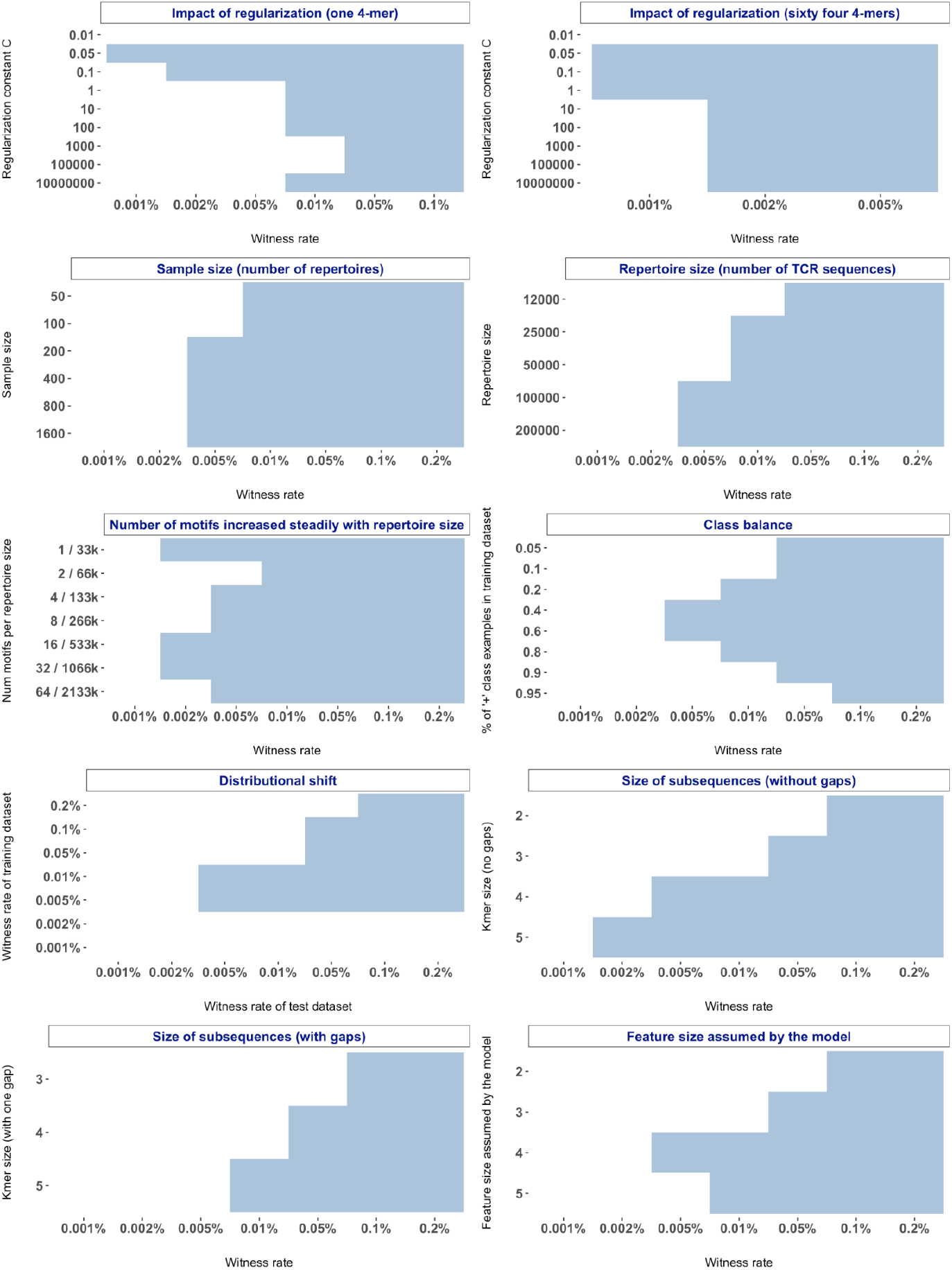
Boundaries and limits of AIRR ML models across variations of dataset and signal properties explored: To demarcate the boundaries where baseline ML methods performed well for AIRR classification and where the baseline methods failed to learn adequately, we discretized the performance profiles of all the investigations (**Figures 2–5)** by a defined threshold of average accuracy ≥ 0.7 and standard deviation ≤ 0.1. The regions in blue show where the baseline methods performed adequately well and the white portions represent scenarios that need particular attention in future benchmarking endeavours of existing AIRR ML methods as well as future methodological developments of sophisticated ML approaches if those scenarios remain intractable for existing AIRR ML methods.

The benchmark dataset suite generated in this study assumes that the signal that distinguishes immune states is embedded within the beta chain of TCRs in the form of linear subsequences (k-mers), which aligns with the observations that that the majority of the possible contacts between TCR and peptide antigens were made through only short and typically linear stretches of amino acid residues of CDR3s (IMGT positions 107–116) [25,33]. Nevertheless, existing AIRR ML methods made varied assumptions regarding the immune signal [18] and reported decent performance metrics on their respective study datasets. For instance, existing methods assumed that immune signal occurs as (a) short linear subsequences (k-mers) [22,38–40], (b) full sequences with/without a particular combination of V gene usage [25,36,37], (c) alternative representations of the sequences such as physicochemical properties of the amino acids [22], (d) relevant features of the full sequences that can be captured as latent variables [27,31], and (e) sequence-independent clonal abundance distributions captured as diversity profiles [52]. A comprehensive benchmarking of the state-of-the-art AIRR ML methods thus requires an even more specialized benchmark dataset suite than those generated in this study to assess the robustness of methods across varying assumptions of the ground truth immune signal.

We obtained several useful insights through profiling the baseline performance of AIRR ML models in this study that will aid both the users and methodology developers of AIRR ML models. First, we noticed that a traditional ML method like logistic regression even with stronger regularization failed to distinguish the classes by immune states at moderate-low witness rates (≤ 0.01%) when the total number of phenotype-associated sequence patterns is large (of which a fraction occurs in positive class examples) (**Figure 2.b)**. Some of the existing AIRR ML methods [22,27,31] aptly treated the repertoire classification problem as a two-staged approach known as multiple instance learning (MIL), where the phenotype-associated patterns are first determined followed by predicting a repertoire-level label based on some form of pooling function of the phenotype-associated patterns. Future studies should investigate if existing MIL-based methods or novel MIL approaches would perform well at moderate-low witness rates (≤ 0.01%) when the total number of phenotype-associated sequence patterns is large.

Second, we found that in our experimental setup, small sample sizes (≤100) were under-powered to learn rare immune signals in this study and a marked improvement in performance was observed by increasing the sample size to 200. On the other hand, despite an 8-fold increase in sample size from 200 repertoires to 1600 repertoires, the improvement in performance of the tested models was not substantial at lower witness rates (≤ 0.005%) (**Figure 3.a**). These observations have to be interpreted in the context of the constraints of contemporary AIRR ML studies in generating datasets with large sample sizes. Several of the AIRR ML studies [22,24,26,37–40] that particularly generated AIRR datasets on clinical samples are constrained by sample sizes and thus trained ML models on sample sizes ≈ 50 repertoires, which is an under-powered sample size to learn signals at low witness rates in this study. A very few contemporary studies have generated datasets with as large sample size as explored in this study. For instance, the recent ImmuneCODE database [35] containing SARS-CoV-2-specific TCR datasets has a curated sample size of ≈ 1500 repertoires. Overall, these observations indicate that if the assumptions of the immune signals used in the benchmark datasets of this study matches the ground truth, then the sample sizes of AIRR ML studies may have to be substantially increased (beyond what is explored in this study) to be able to observe a considerable improvement in the performance of ML models at lower witness rates.

Third, our findings demonstrate that increasing the sequencing depth is more beneficial for training a generalizable model (than increasing sample size) as it increases the likelihood to include and learn more phenotype-associated sequence patterns, especially at moderate-low witness rates (≤ 0.01%) (**Figure 3**). An increase in repertoire size not only improved the performance of the model at moderate-low witness rates (**Figure 3.b**), but also improved the performance when the total number of phenotype-associated sequence patterns is large (**Figure 3.c**). However, it might be a challenge for current experimental datasets to achieve as high sequencing depth as 10^6^ unique sequences on average for all the repertoires and is currently limited to sizes around 10^5^ unique sequences on average with a large dispersion [25,35]. Future studies should investigate the trade-off between sample size and sequencing depth to understand if an increase in sample size would compensate for the shallow sequencing depths.

Although class imbalance [53] and distributional shifts [47] are known phenomena impacting the generalizability of ML models, we here charted out precisely how they impact the generalizability of AIRR ML models (**Figure 4**). We noticed a decrease in the performance of ML models with increased class imbalance. A straightforward approach used often to mitigate the class imbalance effect is to use some form of subsampling to bring balance in the classes. However, such techniques would further reduce the sample size and require a decently large number of examples of the least prevalent class to mitigate the impact of sample size described above. Furthermore, we observed that a distributional shift involving a decrease in witness rate of either the training dataset or test dataset resulted in a decrease of prediction accuracy although the impact of the latter was substantial. While the users of AIRR ML methods should be mindful of the class imbalances and potential distributional shifts, the AIRR ML community should explore, understand and develop tailored approaches that remain robust to class imbalances and distributional shifts in AIRR datasets.

Previous studies reported motifs of different sizes contributing to the epitope specificity ranging from a single amino acid residue [54] to full sequences [25]. In the light of such observations, it is important for the ML models to remain robust irrespective of the size of the ground truth k-mers. However, in this study, we noticed a marked deterioration of prediction performance when the assumed feature size did not match the ground truth, where a much higher witness rate was needed to compensate for a mismatched modeling assumption. The users of AIRR ML methods should thus notice that assuming the ground truth signal as a single fixed feature size can turn out to be a major limitation of the models. Methods that learn relevant features of the sequences as latent variables [27,28,31] may overcome such limitations and thus should be investigated in future benchmarking endeavours. Further, we also observed that shorter k-mers were much harder to learn than longer k-mers, and that complex k-mers (i.e. containing a gap) were much harder to learn than contiguous k-mers. Future studies should investigate if the state-of-the-art AIRR ML methods are able to perform better in learning shorter and complex k-mers and develop methodologies to overcome these challenges faced by traditional encodings and ML methods.

Our exploration of the noise in negative class can be viewed as a case of class noise [55] that sheds light on the signal frequency distributions, where the classes become inseparable.We observed that a model was able to perform well even when the negative class examples carried noise at a rate equivalent to half the witness rate in the positive class examples. In other words, when positive class repertoires carried signal on average in 10 sequences out of 100’000 sequences, the model was able to separate them from negative class repertoires with decent accuracy as long as the negative class repertoires did not carry signal in 5 out of 100’000 sequences. In this simulation study, we were able to assess at what frequency level the ground truth signals are occurring in the negative class as we know the ground truth signals from the outset. When the ground truth signals are not known in experimental datasets, comparing the feature frequency distributions of positive and negative class examples for top features (selected based on model coefficients or some form of feature importance) can act as model diagnostic.

In this study, we generated the synthetic reference repertoires (prior to simulating immune state-associated signal) according to the VDJ recombination model provided by OLGA [42], which acted as a reasonable proxy for real-world experimental repertoires (Figure S8). Although the synthetic repertoires used in this study retain the nativeness of AIRR datasets in terms of positional biases, amino acid usage, sequence length distributions and typical repertoire sizes, future studies should also explore and understand how to mimic the other noises and biases that are inherent to experimental datasets (e.g., sequencing artifacts, library preparation issues, batch effects and impact of other covariates, co-existence of other signals that can further increase the complexity and so on). A natural extension would then be to devise analytical strategies and ML methods that not only handles the idiosyncrasies of experimental datasets but also accounts for the effect of covariates to establish true causal relations between sequence patterns and immune states. Furthermore, the strategy of implanting k-mers as a means to introduce immune-state associated signal into the repertoires was found to be adequate as the implanted k-mers did not induce significant changes in the underlying baseline frequency distributions of other k-mers, thus not disrupting the positional biases of immune receptor sequences substantially (Figure S1). In future work, the impact of k-mer implantation on TCR generation probabilities should be investigated in order to investigate to what extent motif implantation changes the a priori likelihood of a sequence [4,56]. In this study, we performed all the experiments using the TCRβ CDR3 amino acid sequences. However, the findings are applicable to adaptive immune receptor repertoires in general given that BCR and TCR sequences have very similar immunogenic architecture. That said, it will be of interest to measure the impact of BCR somatic hypermutation in the (BCR) AIRR ML applications [41,57].

The importance of the technical implementation setup of a large scale computational study such as the one carried out here is noteworthy. To carry out a similar study involving AIRR ML models, one would need streamlined operations to (a) read-in AIRR-seq datasets in different file formats (both experimental and synthetic), (b) represent the information of datasets in multiple ways (encoding schemes), (c) simulate immune state-associated signals, (d) have access to a wide variety of ML model implementations, (e) perform model interpretation easily through a range of exploratory diagnostic analyses, (f) ensure reproducibility, shareability, and transparency of the analyses. Handling all the aforementioned operations even when building on top of existing ML frameworks such as scikit-learn [46] with *ad hoc* scripts can be challenging, time-consuming (for all the boilerplate code implementations), and less efficient if the implementations are not well-tested or optimized. To overcome all these challenges, we used immuneML [58], which is a domain-adapted software ecosystem for ML analysis of AIRR datasets.

## Conclusions

To help the scientific community in avoiding futile efforts of developing novel ML methodology for those scenarios of AIRR classification, where baseline methods may already excel, we profiled the baseline performance of AIRR ML models across a diverse set of challenges with increasing levels of complexities for the classification problem. The summarized findings (**Figure 6**) showed the boundaries in terms of AIRR dataset and immune signal characteristics, where baseline ML methods are able to classify repertoires according to immune state. Future benchmarking studies should investigate if the state-of-the-art AIRR ML methods are able to perform well on the intractable scenarios identified in this study. Such an endeavour can help narrow down the scenarios of AIRR classification where novel methodology development is needed. The benchmark dataset suite and the knowledge on baseline performance levels of AIRR ML models generated in this study serve multiple purposes by: (a) providing a reference benchmark for novel AIRR classification methodology and enumerating scenarios where novel methodology development is not needed. (b) providing realistic expectations of the performance of AIRR ML models given the training dataset properties and assumptions. (c) serving as a template for defining specialized AIRR benchmark datasets for comprehensive benchmarking of the AIRR ML methods.

## Methods

### Generation of synthetic AIRR reference datasets

In this study, our goal was to profile the performance of AIRR-ML models across a wide range of distinct challenges listed in **Table 1**. Ideally, such an endeavour is best carried out with a combination of experimental and simulated datasets. However, often the ground truth in experimental datasets is not known. The impact of different variations of AIRR dataset properties and signal properties on the performance of ML models cannot be sufficiently teased apart if there exists no knowledge on how hard the learning problem is at the outset. Therefore, we chose to generate synthetic AIRR reference datasets that retain the realistic nature of AIRR datasets in terms of positional biases, amino acid usage in the sequences of repertoires, sequence length distributions and typical repertoire sizes. We generated the desired number of repertoires (default 200) for each benchmark dataset by generating a desired number of TCRβ CDR3 amino acid sequences (default 100’000) according to the VDJ recombination model provided by OLGA [42]. In one of the benchmarks, we sought to evaluate whether the performance of ML models would vary if the datasets used for training the models deviate from AIRR-specific characteristics. For that, in addition to the repertoires generated according to the OLGA-provided VDJ recombination model, we generated repertoire datasets in three other ways: (a) desired number of repertoires composed of desired number of random amino acid sequences matching the length distribution of typical CDR3 amino acid sequences. (b) desired number of repertoires composed of desired number of amino acid sequences randomly sampled from a large pool of experimental TCRβ CDR3 amino acid sequences from Emerson et al [25]. (c) desired number of whole repertoires from Emerson et al [25] that have comparable repertoire sizes achieved through subsampling. Hereafter we refer to the four synthetic reference datasets generated here as OLGA-based repertoires, random repertoires, meta repertoires (the one with pooled sequences from experimental dataset), and experimental repertoires (subsampled to have comparable repertoire sizes).

### Construction of benchmark datasets through the simulation of immune state-associated signals

In all except one benchmark dataset mentioned above (Table 1), we used OLGA-based repertoires to construct the benchmark datasets. Existing knowledge suggests that the majority of the possible contacts between TCR and peptide antigens were made through only short and typically linear stretches of amino acid residues of CDR3s (IMGT positions 107–116) [25,33]. To account for the often unknown and varying lengths of the subsequences that make contact with peptide antigens, some of the previous studies either used a fixed assumption regarding the subsequence size (e.g., 4-mers [22]) or used varied definitions of the subsequence size (e.g., 2-mers, 3-mers, 4-mers and 5-mers [20,26]). In this study, while we use varying definitions of k-mer size to assess the impact of signal definition on classification performance, we use a fixed definition (4-mers) when the aims were to assess the impact of other properties of the AIRR ML training setup. As a default definition, we used a signal composed of three k-mers of size 4 (4-mers) in the vast majority of the benchmarks. In each of the reference dataset, signal is implanted at multiple different witness rates (0.001%, 0.002%, 0.005%, 0.01%, 0.05%, 0.1%, 0.2%) in 50% of the dataset. The signal was implanted with equal probability at IMGT sequence positions 108–111 to not disrupt the germline signal in the conserved positions (the start and end portions of the sequences). The examples in which the signal was implanted were labeled positive class (with a hypothetical immune state) and the remaining were labeled as negative class. In addition, for each investigation, one of the properties pertaining to the dataset, signal or ML model was varied at a range of values. **Table 1** shows a list of the different dataset or signal properties that were varied for the construction of benchmark datasets and the particular variations (range of values) of each property that were explored. For each property that was varied, a benchmark dataset was constructed at all the witness rates explored. When investigating the noise in the negative class, the signal was implanted in the positive class examples at only one fixed witness rate (0.1%). **Figure 1.b** shows a schematic illustration of implanting an immune state-associated sequence signal into synthetic AIRR reference datasets, and **Figure 1.c** shows schematic illustrations of the construction of benchmark datasets.

### Sequence encoding and preprocessing

A k-mer frequency encoding was used in all the ML model training setups of this study. In other words, all the ML models of this study assume that the immune signal that differentiates positive and negative class occurs in the form of a contiguous subsequence of a defined size ***k*** (k-mers). The sequences in each repertoire were split into overlapping k-mers and their frequencies were computed followed by L2 normalization of k-mer frequencies. Furthermore, each feature vector was standardized by scaling variance to one and mean to zero across examples. The default encoding scheme was subsequences of size 4 (4-mers). In a subset of the benchmarking experiments, the ***k*** in k-mer frequency encoding varied from 2 to 5.

### Machine learning models

We used Scikit-learn’s [46] implementation of a *L1* regularized logistic regression (lasso [59]) for the large majority of the benchmarking experiments in order to establish a baseline for the predictive performance of ML models across diverse levels of challenges of AIRR classification. Consider a response variable *Y* with two classes ‘+’ and ‘-’ and predictor variables *X* with *X*_*i*_ denoting the ith example and *X*_*ij*_ denoting the jth feature of the ith example. In logistic regression, the conditional probability of P(*Y*=+|*X*), shortly p(*X*), is modeled using the logit function 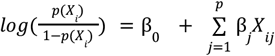 and the values of β_0_ and β_j_ are found through minimising the negative log likelihood given by the equation: 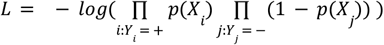 where П represents product over *i* and *j* that run over positive (+) and negative (-) classes respectively. In *L1* penalized logistic regression, the cost function that is minimised is 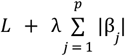, where p is the number of predictors. Note that the hyperparameter controlling regularization strength is indicated by a variable **C** in Scikit-learn’s [46] implementations and is the inverse of regularization strength (C = λ^-1^). Smaller values of C represent stronger regularization. The hyperparameter space used for controlling the regularization strength constant **C** was by default 0.05, 0.1, 1, 5 (except for the benchmarking experiment that specifically explored the effect of regularization, where **C** varied across a wider range of values). A maximum of 500 iterations were allowed for the model’s coefficients to converge. Although a comprehensive assessment and benchmarking of ML methods for AIRR classification falls outside the scope of this study, we briefly explored how straightforward application of other classical ML methods like Random Forests (RF) or Support Vector Classifier (SVC) compared to the logistic regression model tested throughout this manuscript. The hyperparameter space for RF and SVC were narrowed down in a similar fashion as logistic regression. The regularization constant space and the maximum number of iterations used for SVC were the same as logistic regression. For RF, the number of trees explored in hyperparameter optimization were 5, 10, 50 and 100.

### Model training, selection and evaluation

To estimate the prediction performance of the trained ML models on unseen AIRR data, we used a nested cross-validation (CV) setup. Briefly, we used a 5-fold nested CV, where in the outer loop 80% of the data is used for training the model and the remaining 20% as a test set for estimating the model performance. In the inner loop of the nested CV, we use a 5-fold CV where the training data is again further split into 80% training and 20% validation set for tuning the hyperparameters and aid in model selection. An exhaustive grid search was used for tuning the hyperparameters. The performance metric optimized in model training was accuracy. Note that we used balanced datasets for all the benchmarking experiments, except one scenario where the impact of class balance was explored.

### Assessment of the impact of k-mer implantation on background k-mer frequency distribution

We assessed the impact of k-mer implantation on background k-mer frequency distributions in two different scenarios: (a) one where three 4-mers are implanted and (b) another where 64 4-mers are implanted. We used 200 repertoires generated by OLGA with a repertoire size of 100’000 sequences. We made an assumption that the 4-mer frequencies of the 200 repertoires will not differ in the absence of any immune state-associated sequence pattern (before 4-mer implantation). Thus, if we would compare the k-mer frequency distributions of two groups (the repertoires that would later become positive and negative class labels after implantation), there should be no significant differences between the groups. For this, we used a student’s t-test to compare the two group frequency distributions for each k-mer (as many number of tests as the number of k-mers) followed by a multiple testing correction using Benjamini-Hochberg method. We repeated the same process of statistical testing on datasets that received implantation of 4-mers at different witness rates in each scenario (three 4-mers implanted or 64 4-mers implanted). We thresholded on the multiple testing corrected p-values (q-value<0.05) to identify 4-mers that exhibited differential incidence. The experiments were replicated on three separate datasets and the average and standard deviation of the number of differentially incident motifs at different witness rates were reported.

### Implementation details

A large majority of the steps of this benchmarking study were carried out using immuneML [58] (version 1.0.2), an integrated ecosystem for machine learning analysis of adaptive immune receptor repertoires. The ML methods used in this study were based on Scikit-learn’s [46] implementations. Specifically, the simulation of disease signals into the AIRR datasets for the construction of benchmark datasets, the sequence encoding and preprocessing steps, and the model training, selection and evaluation were performed through immuneML.

### Docker container to improve reproducibility

To enable other researchers to re-use and reproduce the findings of this manuscript, we created a Docker [60] image with a predefined computing environment maintaining all the dependencies required for execution of the code with minimal overhead. Although newer versions of immuneML [58] and its dependencies have emerged during the preparation of this manuscript, the Docker image froze the exact versions of all the software and thereby eliminates any barriers of reproducing the findings of this study. The Docker image can be accessed from the publicly hosted central repository of Docker (dockerhub): *kanduric/immuneml-v1:latest*. Additionally, we demonstrated the ease of re-usability and deployability of the containerized computational workflow by re-running a subset of analyses from each experiment of this study on a toy dataset (100 sequences in each repertoire compared to 100’000 in the original analyses) with fewer iterations in ML training [61]. Although the findings of such analyses are expected to be illogical, the main idea behind choosing a toy dataset and fewer iterations in ML training for this demonstration purpose is to ensure rapid testing by other researchers on different computational environments. Notably, the original computational analysis of this manuscript involves reading in ∼ 2 TB of data from the disk and writing ∼10 TB of data to the disk. To replicate the findings of this manuscript, the original input data [62], analysis scripts [63], and the docker image that are made publicly available have to be used in a similar fashion as shown in the demo analyses [61].

### Graphics

ggplot2 [64] was used for graphs and inkscape [65] was used for illustrations.

## Data and Source Code availability

The bash scripts used to generate all the synthetic repertoires are made publicly available. All the synthetic repertoires (∼ 2.1 TB) used as input for the simulation of disease signals and ML model training are made publicly available: https://doi.org/10.11582/2021.00064. The analytical details for all investigations including simulation of disease signals and ML model training are made available in the form of immuneML YAML specification files that describe each step and choice of the analyses: https://doi.org/10.11582/2021.00038. These specifications are both human-readable (analysis transparency) and executable using the publicly available immuneML platform [58] version 1.0.2. To ensure the ease of re-usability and reproducibility, we containerized the whole computational environment that is necessary to remove any barriers of replicating the findings of this study. The docker image of the computational workflow of this manuscript is available from publicly hosted docker repository (dockerhub) at *kanduric/immuneml-v1:latest*. A demo of using the provided docker image to re-run each category of experiment in this manuscript has been shown on toy datasets at: https://github.com/KanduriC/demo_reproducibility_kanduricetal2021.git

## Declarations

### List of abbreviations

ML: Machine Learning
AIR: Adaptive Immune Receptors
AIRR: Adaptive Immune Receptor Repertoires
TCR: T Cell Receptors
TCRβ: T Cell Receptor beta chain
CDR3: Complementarity Determining Region 3
SVC: Support Vector Classifier
RF: Random Forests
CV: Cross Validation
IMGT: ImMunoGeneTics

## Ethics approval and Consent

Not applicable

## Consent for publication

Not applicable

## Authors’ contributions

CK and GKS conceived the overall study. MP, LS, KM, MC and VG participated in brainstorming and provided critical conceptual feedback. CK performed all the analyses and drafted the manuscript. VG and GKS added critical edits to the manuscript. All authors read and approved the final manuscript.

## Acknowledgements

This work was performed using the Immunohub eInfrastructure funded by University of Oslo and operated by the authors in close collaboration with the University Senter for Information Technology (USIT), University of Oslo.

---

**Table S1:**
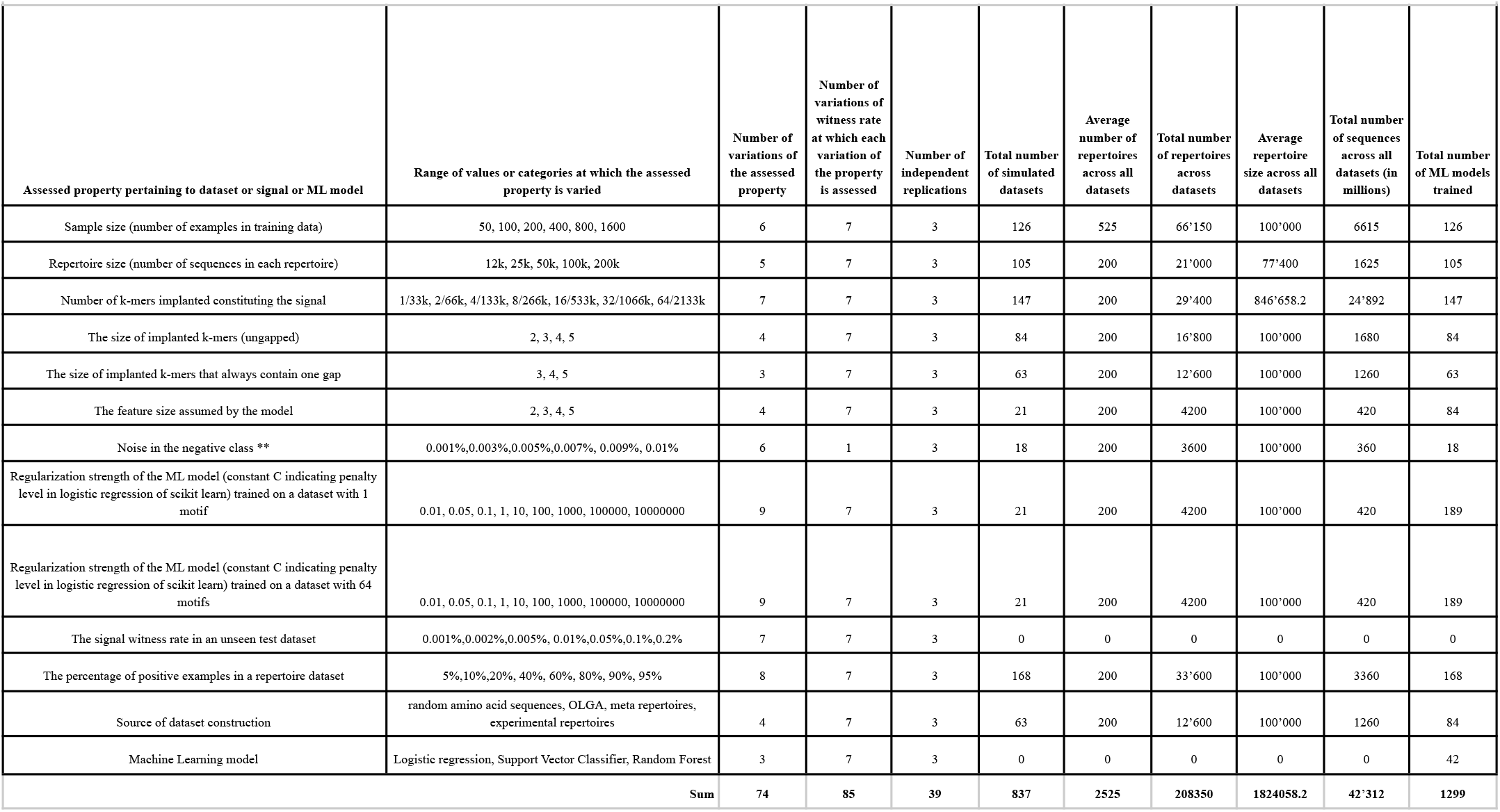
Descriptive statistics of the benchmarking setup.

| Assessed property pertaining to dataset or signal or ML model | Range of values or categories at which the assessed property is varied | Number of variations of the assessed property | Number of variations of witness rate at which each variation of the property is assessed | Number of independent replications | Total number of simulated datasets | Average number of repertoires across all datasets | Total number of repertoires across datasets | Average repertoire size across all datasets | Total number of sequences across all datasets (in millions) | Total number of ML models trained |
| --- | --- | --- | --- | --- | --- | --- | --- | --- | --- | --- |
| Sample size (number of examples in training data) | 50, 100, 200, 400, 800, 1600 | 6 | 7 | 3 | 126 | 525 | 66'150 | 100'000 | 6615 | 126 |
| Repertoire size (number of sequences in each repertoire) | 12k, 25k, 50k, 100k, 200k | 5 | 7 | 3 | 105 | 200 | 21'000 | 77'400 | 1625 | 105 |
| Number of k-mers implanted constituting the signal | 1/33k, 2/66k, 4/133k, 8/266k, 16/533k, 32/1066k, 64/2133k | 7 | 7 | 3 | 147 | 200 | 29'400 | 846'658.2 | 24'892 | 147 |
| The size of implanted k-mers (ungapped) | 2, 3, 4, 5 | 4 | 7 | 3 | 84 | 200 | 16'800 | 100'000 | 1680 | 84 |
| The size of implanted k-mers that always contain one gap | 3, 4, 5 | 3 | 7 | 3 | 63 | 200 | 12'600 | 100'000 | 1260 | 63 |
| The feature size assumed by the model | 2, 3, 4, 5 | 4 | 7 | 3 | 21 | 200 | 4200 | 100'000 | 420 | 84 |
| Noise in the negative class ** | 0.001%,0.003%,0.005%,0.007%, 0.009%, 0.01% | 6 | 1 | 3 | 18 | 200 | 3600 | 100'000 | 360 | 18 |
| Regularization strength of the ML model (constant C indicating penalty level in logistic regression of scikit learn) trained on a dataset with 1 motif | 0.01, 0.05, 0.1, 1, 10, 100, 1000, 100000, 10000000 | 9 | 7 | 3 | 21 | 200 | 4200 | 100'000 | 420 | 189 |
| Regularization strength of the ML model (constant C indicating penalty level in logistic regression of scikit learn) trained on a dataset with 64 motifs | 0.01, 0.05, 0.1, 1, 10, 100, 1000, 100000, 10000000 | 9 | 7 | 3 | 21 | 200 | 4200 | 100'000 | 420 | 189 |
| The signal witness rate in an unseen test dataset | 0.001%,0.002%,0.005%, 0.01%,0.05%,0.1%,0.2% | 7 | 7 | 3 | 0 | 0 | 0 | 0 | 0 | 0 |
| The percentage of positive examples in a repertoire dataset | 5%,10%,20%, 40%, 60%, 80%, 90%, 95% | 8 | 7 | 3 | 168 | 200 | 33'600 | 100'000 | 3360 | 168 |
| Source of dataset construction | random amino acid sequences, OLGA, meta repertoires, experimental repertoires | 4 | 7 | 3 | 63 | 200 | 12'600 | 100'000 | 1260 | 84 |
| Machine Learning model | Logistic regression, Support Vector Classifier, Random Forest | 3 | 7 | 3 | 0 | 0 | 0 | 0 | 0 | 42 |
| Sum |  | 74 | 85 | 39 | 837 | 2525 | 208350 | 1824058.2 | 42'312 | 1299 |

## Supplementary Information

**Figure S1.**
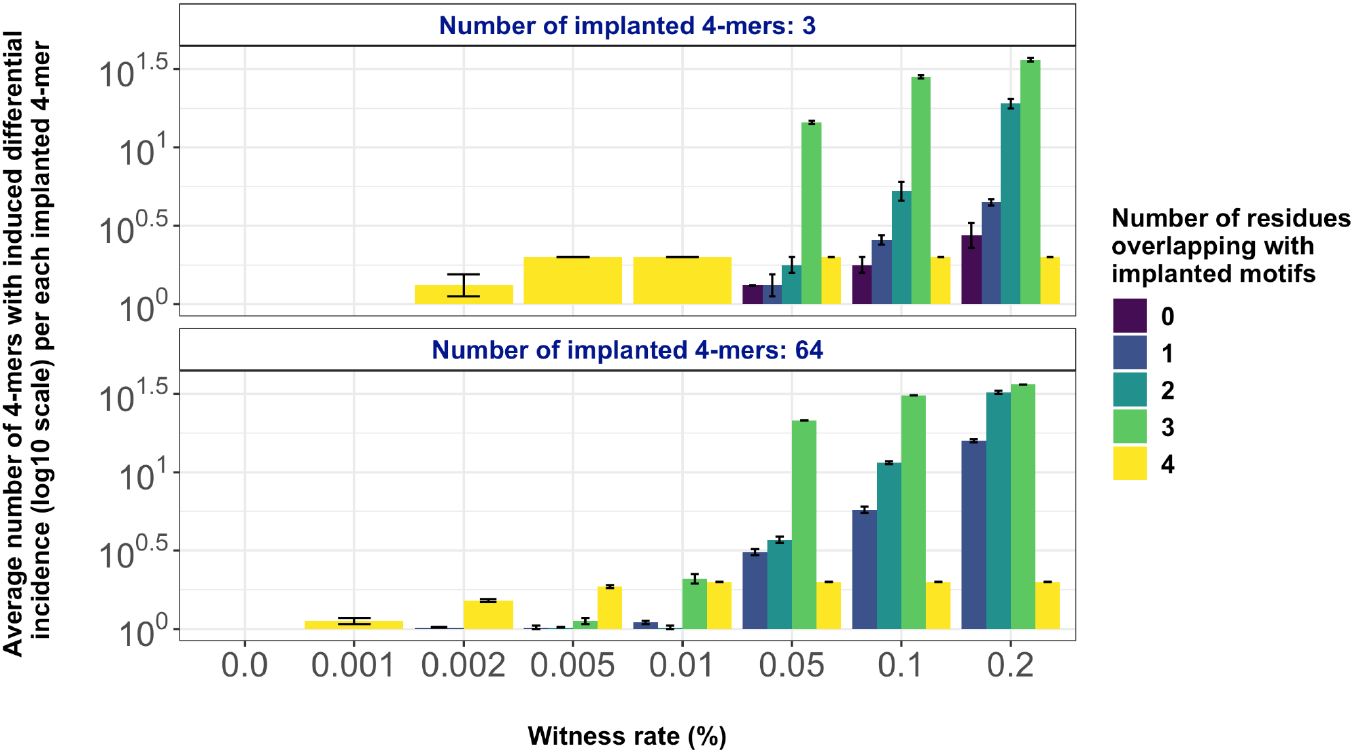
(Relates to Figure 2.b and Figure 3.a). Impact of k-mer implantation on the background k-mer frequency distribution: The y-axis shows the average number of motifs with induced significant differential incidence scaled by the total number of implanted motifs in log_10_ scale when the signal (number of implanted 4-mers in upper and lower panels) was implanted at different witness rates (on x-axis) in 200 repertoires of size 100’000 sequences. The color coding separates the motifs with induced differential incidence by how much they overlap with any implanted motif. For instance, the yellow color indicates that the differentially incident motif overlaps all its 4 amino acid residues with one of the implanted 4-mers. In other words, those are the true implanted motifs. The error bars are based on the standard deviation of values across three independent replications. The chart shows that at lower witness rates (up to 0.01%), the implantation of 4-mers predominantly disturbs the frequency distributions of only the implanted 4-mers between positive and negative class repertoires. As the witness rate increases, the signal gets implanted in more sequences thus overlapping with many different 4-mers by 1, 2 or 3 amino acid residues. The total number of disturbed 4-mers increased proportionally to the number of implanted 4-mers (as evident through the scaling of y-axis per the number of implanted 4-mers).

**Figure S2.**
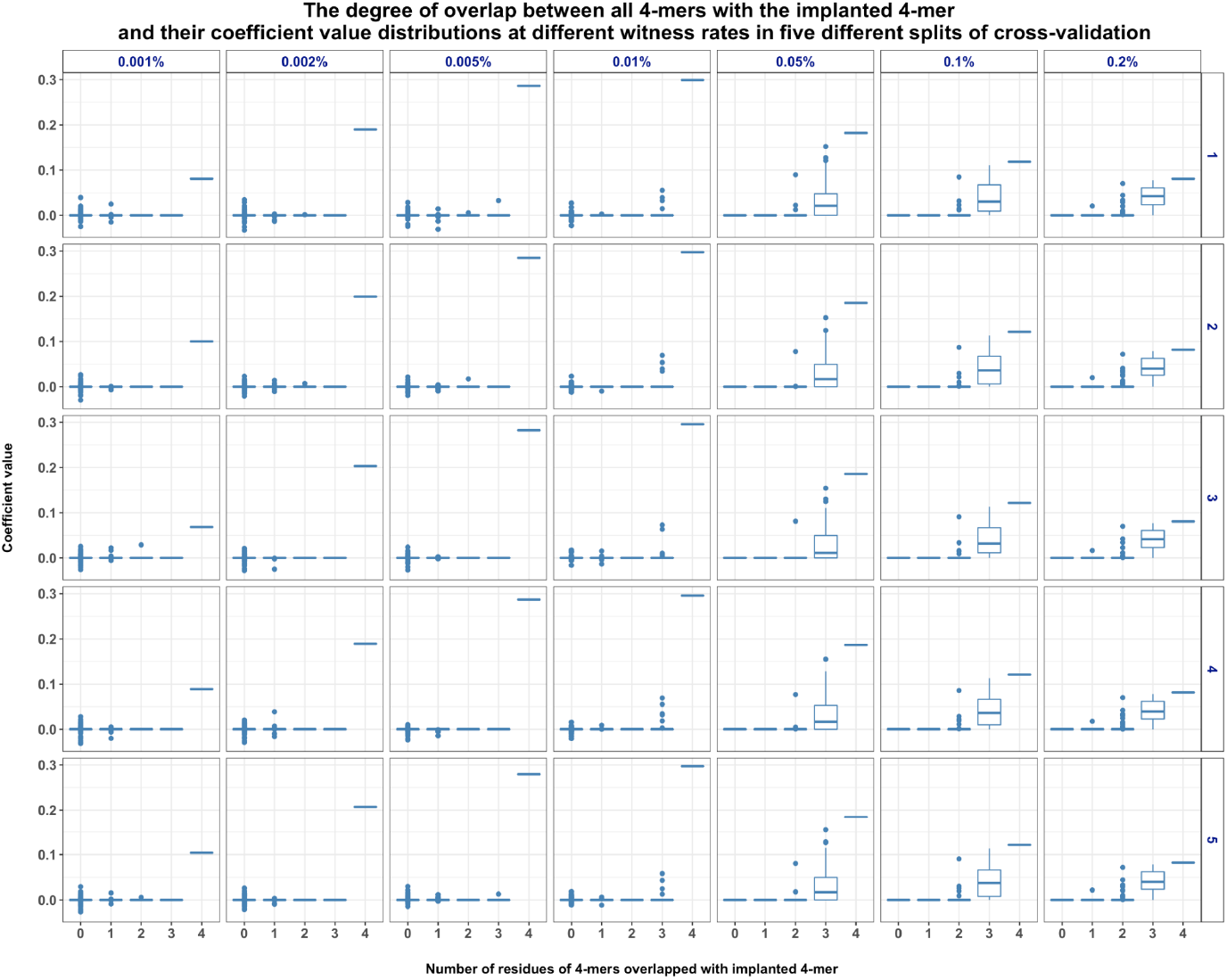
(Relates to Figure 2.a). Distribution of the coefficient values of the trained model categorized by the level of overlap with the implanted single 4-mer (as in Figure 2.a). The y-axis shows the coefficient values of different 4-mers and the x-axis shows the number of amino acid residues of the 4-mers that overlap the true implanted motifs. In this case, an overlap value of 4 indicates that the 4-mer is the true implanted motif. Different panels in columns show different witness rates and the panels in different rows correspond to the different splits of the cross-validation. The charts confirm that at lower witness rates, it was only the truly implanted motif that was ascribed the maximum weight (coefficient) by the model. As the witness rate increases, the motif gets implanted in more sequences and thereby overlaps with many other motifs partially by either 1, 2, or 3 amino acid residues. Thus such partially overlapping motifs would also be ascribed higher weights by the model as the witness rate increases.

**Figure S3.**
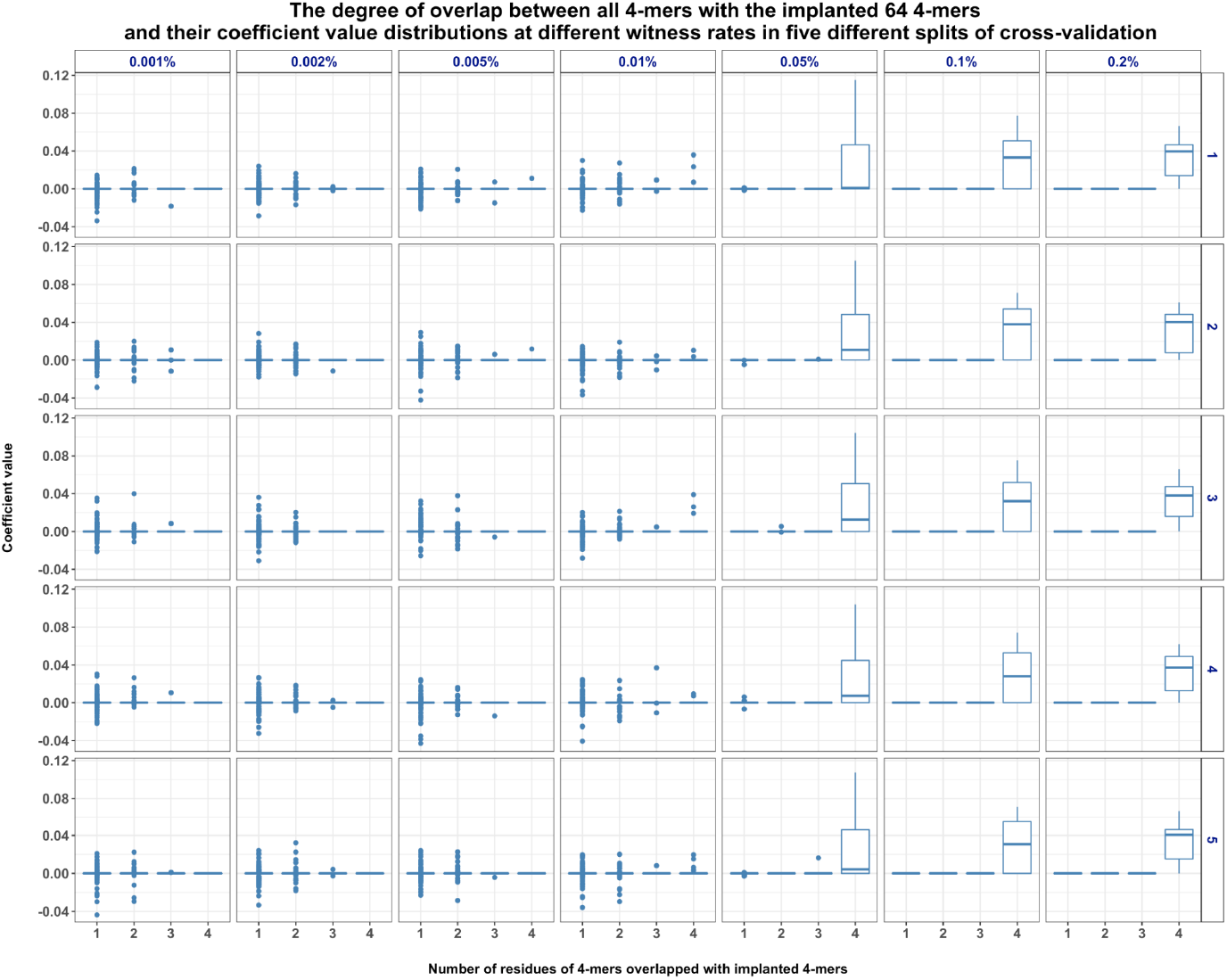
(Relates to Figure 2.b). Distribution of the coefficient values of the trained model categorized by the level of overlap with the implanted 64 4-mers (as in Figure 2.b). The y-axis shows the coefficient values of different 4-mers and the x-axis shows the number of amino acid residues of the 4-mers that overlap the true implanted motifs. In this case, an overlap value of 4 indicates that the 4-mers are the true implanted motifs. Different panels in columns show different witness rates and the panels in different rows correspond to the different splits of the cross-validation. The charts show that at lower witness rates, none of the 4-mers were ascribed extreme weights by the model. As the witness rate increased, the true motifs were ascribed higher weights.

**Figure S4.**
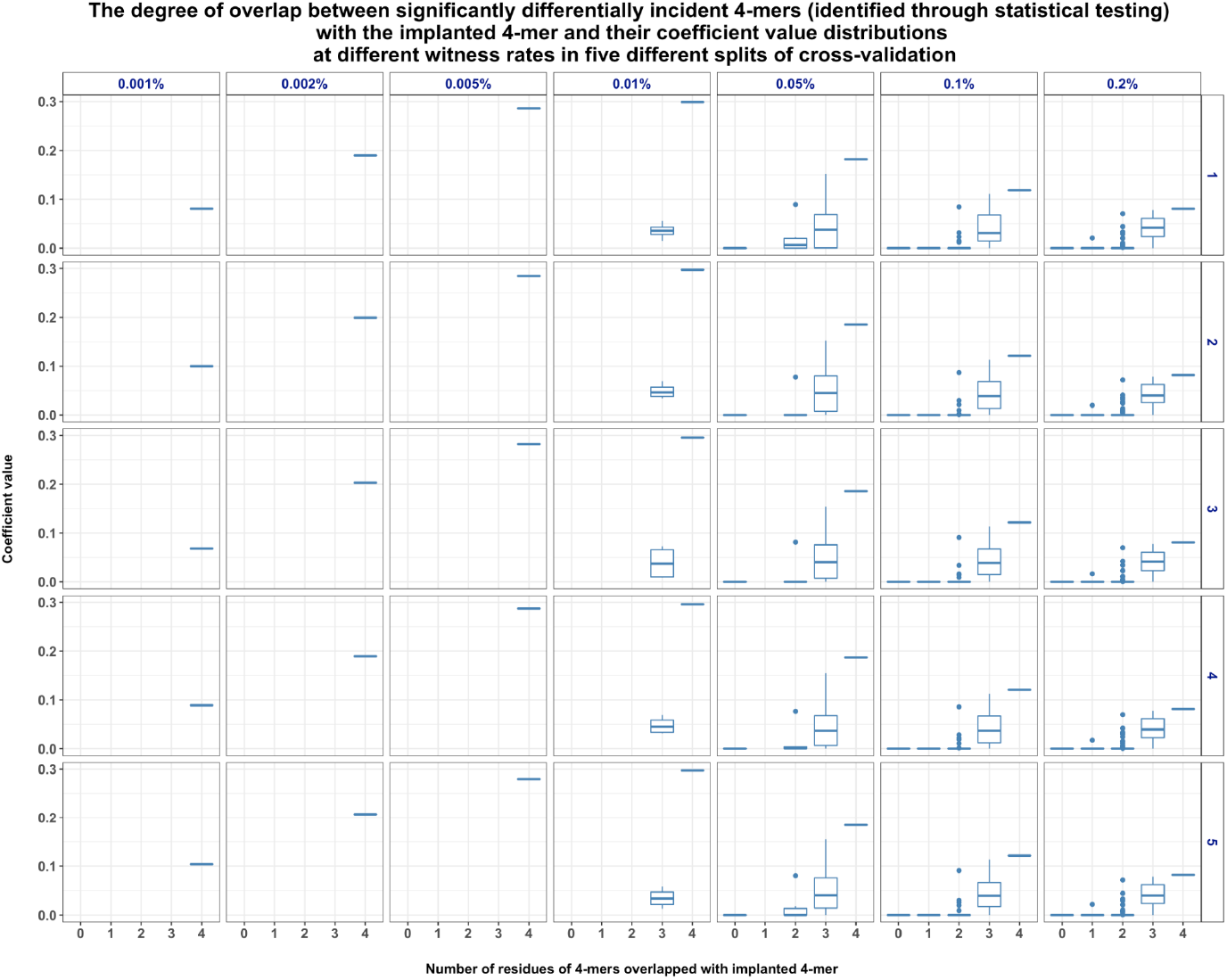
(Relates to Figure 2.a and Figure S2). Distribution of the coefficient values of the trained model categorized by the level of overlap with the implanted single 4-mer (as in Figure 2.a) - this chart unlike Figure S2 shows the distribution only for the statistically significant 4-mers that were identified through a univariate statistical testing. From all the 4-mers that were plotted in Figure S2, we filtered out all those 4-mers that were found with no significant differences in frequency distributions in a univariate statistical test (see Methods) and retained only the significant motifs. This chart confirms that statistically significant motifs that can be identified through a univariate statistical test comparing the frequency distributions are indeed the same motifs that were ascribed higher weights by the penalized logistic regression model. The y-axis shows the coefficient values of different 4-mers and the x-axis shows the number of amino acid residues of the 4-mers that overlap the true implanted motifs. In this case, an overlap value of 4 indicates that the 4-mer is the true implanted motif. Different panels in columns show different witness rates and the panels in different rows correspond to the different splits of the cross-validation. Our analysis confirms that at lower witness rates, it was only the truly implanted motif that was ascribed the maximum weight (coefficient) by the model. As the witness rate increases, the motif gets implanted in more sequences and thereby overlaps with many other motifs partially by either 1, 2, or 3 amino acid residues. Thus such partially overlapping motifs would also be ascribed higher weights by the model as the witness rate increases.

**Figure S5.**
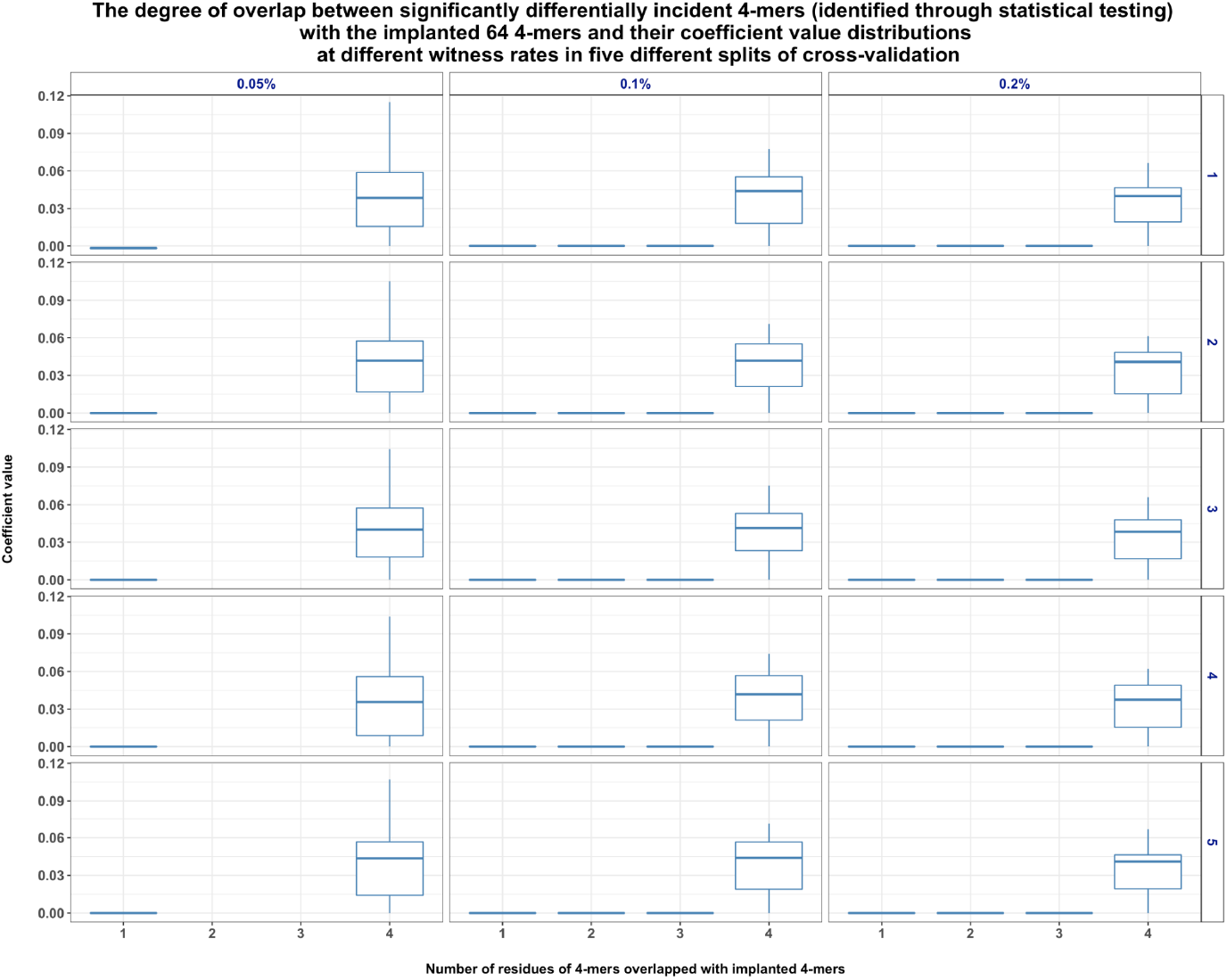
(Relates to Figure 2.b and Figure S3). Distribution of the coefficient values of the trained model categorized by the level of overlap with the implanted 64 4-mers (as in Figure 2.b) – this chart unlike Figure S3 shows the distribution only for the statistically significant 4-mers that were identified through a univariate statistical testing. From all the 4-mers that were plotted in Figure S3, we filtered out all those 4-mers that were found with no significant differences in frequency distributions in a univariate statistical test (see methods) and retained only the significant motifs. This chart confirms that statistically significant motifs that can be identified through a univariate statistical test comparing the frequency distributions are indeed the very same motifs that were ascribed higher weights by the penalized logistic regression model. *Note that unlike Figure S3, this chart shows fewer column panels corresponding to witness rates. This was because there were no significant motifs that were identified at lower witness rates*. The y-axis shows the coefficient values of different 4-mers and the x-axis shows the number of amino acid residues of the 4-mers that overlap the true implanted motifs. In this case, an overlap value of 4 indicates that the 4-mers are the true implanted motifs. Different panels in columns show different witness rates and the panels in different rows correspond to the different splits of the cross-validation. The charts show that at lower witness rates, none of the 4-mers were ascribed extreme weights by the model. As the witness rate increased, the true motifs were ascribed higher weights.

**Figure S6.**
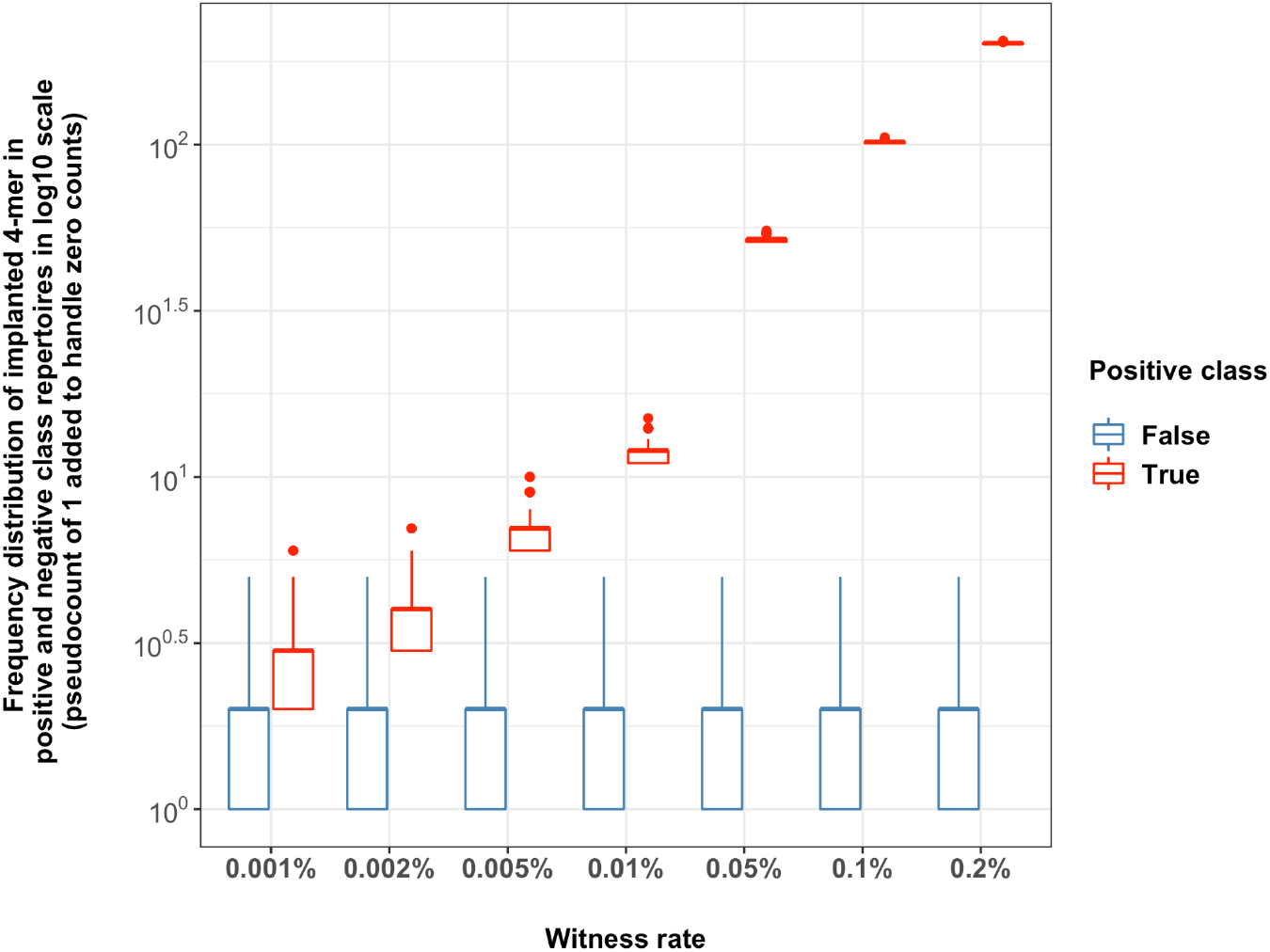
(Relates to Figure 2.a). Frequency distribution of implanted 4-mer (as in Figure 2.a) in both positive and negative class examples. On the y-axis, the frequency of the implanted 4-mer is shown in log10 scale at different witness rates on the x-axis. The color indicates positive and negative class examples. The chart shows that already at a witness rate of 0.002%, the frequency distributions of both positive and negative classes started to become distinguishable.

**Figure S7.**
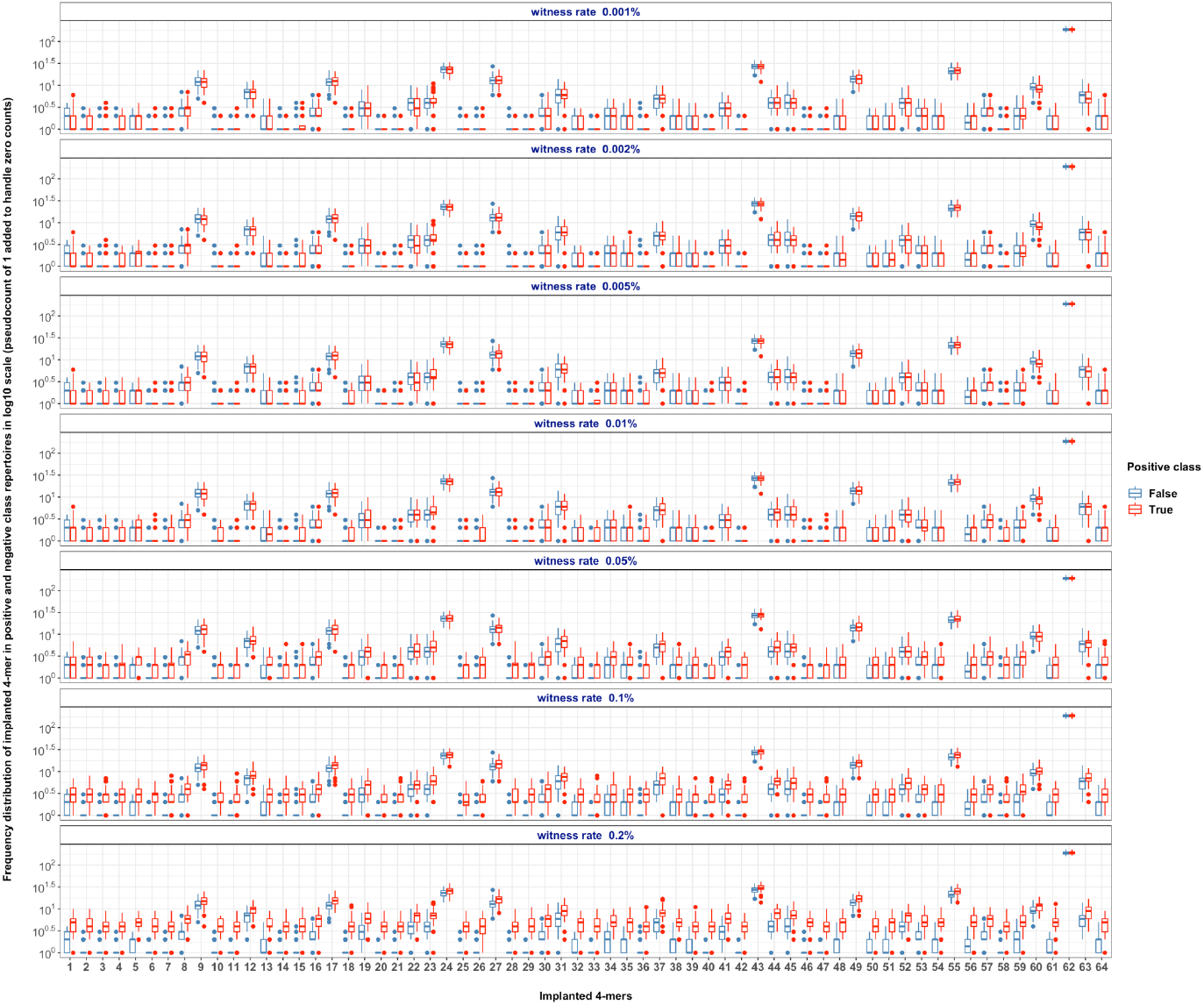
(Relates to Figure 2.b). Frequency distribution of implanted 64 4-mers (as in Figure 2.b) in both positive and negative class examples. On the y-axis, the frequency of the implanted 4-mers is shown in log10 scale at different witness rates (in different panels). The x-axis shows the index of each implanted 4-mer. The color indicates positive and negative class examples. The chart shows that at lower witness rates, the frequency distributions of both positive and negative classes are indistinguishable.

**Figure S8.**
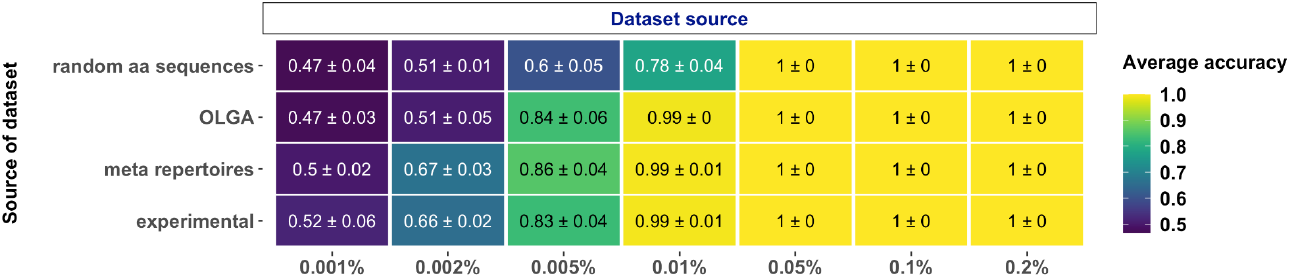
(Relates to Figures 2-5). Impact of the source of dataset construction. Performance estimates of a regularized logistic regression model in a binary classification of a balanced, labeled datasets of varying sources (on y-axis), where the signal in positive class examples composed of 4-mers are known at the explored witness rates (on x-axis).

**Figure S9.**
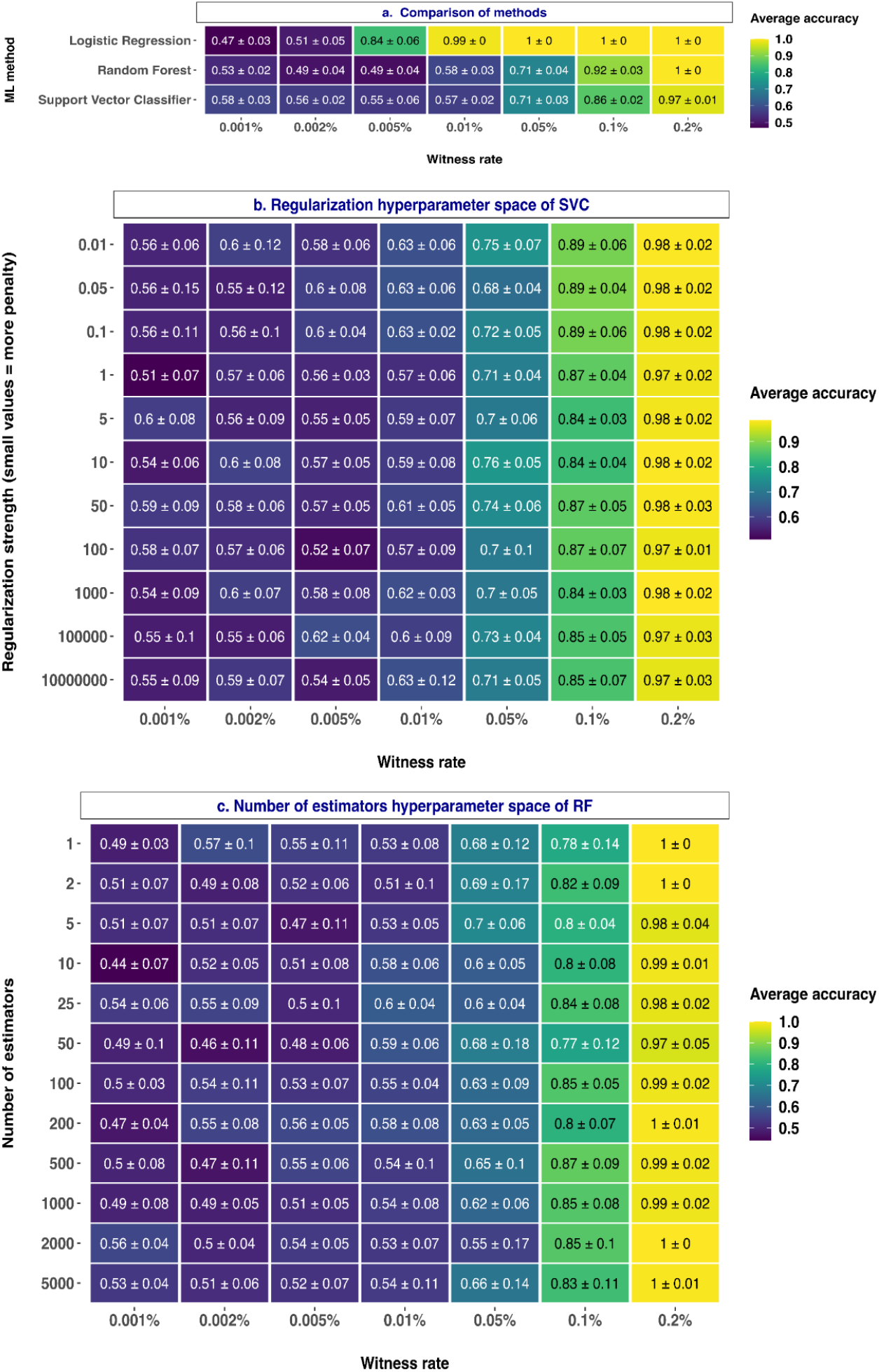
(Relates to Figures 2-5). Impact of the choice of ML method and the hyperparameter spaces of explored ML methods. **(a)** Performance estimates of different ML models (on y-axis) in a binary classification of balanced, labeled AIRR datasets, where the signal in positive class examples composed of 4-mers are known at the explored witness rates (on x-axis). The mean balanced accuracy of a 5-fold cross-validation was computed in three independent replications. The color coding shows the mean and standard deviation of the performance estimate obtained by three independent replications.**(b) Impact of regularization on the performance estimates of SVC**. Performance estimates of a support vector classifier (SVC) regularized with a fixed regularization constant C (explored on y-axis) in a binary classification of a balanced, labeled AIRR dataset where the signal in positive class examples composed of 4-mers are known at the explored witness rates (on x-axis). Smaller the value of regularization constant C, stronger the regularization. The signal definition is composed of three motifs. The color coding shows the mean and standard deviation of the balanced accuracy estimated by a 5-fold cross-validation.**(c) Impact of the number of estimators on the performance estimates of the RF classifier**. Performance estimates of a random forest classifier (RF) parametrized with a fixed number of estimators (explored on y-axis) in a binary classification of a balanced, labeled AIRR dataset where the signal in positive class examples composed of 4-mers are known at the explored witness rates (on x-axis). The signal definition is composed of three motifs. The color coding shows the mean and standard deviation of the balanced accuracy estimated through a 5-fold cross-validation.

